# Modeling reveals the strength of weak interactions in stacked ring assembly

**DOI:** 10.1101/2024.02.02.578706

**Authors:** Leonila Lagunes, Koan Briggs, Paige Martin-Holder, Zaikun Xu, Dustin Maurer, Karim Ghabra, Eric J. Deeds

## Abstract

Cells employ many large macromolecular machines for the execution and regulation of processes that are vital for cell and organismal viability. Interestingly, cells cannot synthesize these machines as functioning units. Instead, cells synthesize the molecular parts that must then assemble into the functional complex. Many important machines, including chaperones like GroEL and proteases like the proteasome, are comprised protein rings that are stacked on top of one another. While there is some experimental data regarding how stacked-ring complexes like the proteasome self-assemble, a comprehensive understanding of the dynamics of stacked ring assembly is currently lacking. Here, we developed a mathematical model of stacked trimer assembly, and performed an analysis of the assembly of the stacked homomeric trimer, which is the simplest stacked ring architecture. We found that stacked rings are particularly susceptible to a form of kinetic trapping that we term “deadlock,” in which the system gets stuck in a state where there are many large intermediates that are not the fully-assembled structure, but that cannot productively react. When interaction affinities are uniformly strong, deadlock severely limits assembly yield. We thus predicted that stacked rings would avoid situations where all interfaces in the structure have high affinity. Analysis of available crystal structures indicated that indeed the majority – if not all – of stacked trimers do not contain uniformly strong interactions. Finally, to better understand the origins of deadlock, we developed a formal pathway analysis and showed that, when all the binding affinities are strong, many of the possible pathways are utilized. In contrast, optimal assembly strategies utilize only a small number of patwhays. Our work suggests that deadlock is a critical factor influencing the evolution of macromolecular machines, and provides general principles for not only understanding existing machines but also for the design of novel structures that can self-assemble efficiently.

**Statement of Significance:** Understanding the assembly macromolecular machines is important for understanding a wide range of cellular processes. Here, we use mathematical models to study the assembly of stacked rings, which are a common motif in these machines. Our models revealed that these complexes can readily get “stuck” during assembly when the binding affinity between subunits is too strong. This suggests an evolutionary pressure to favor weaker interactions, and our analysis of solved structures confirmed this prediction. Our findings not only contribute to the fundamental understanding of assembly but also offer insights into the evolutionary pressures shaping the architecture of stacked rings, and have implications for both cell and synthetic biology.

## Introduction

Cells employ large macromolecular machines for the execution and regulation of almost every process vital to cell and organismal viability. For instance, protein homeostasis relies on the ribosome for protein synthesis (1), the proteasome for protein degradation (2) and chaperones like GroEL to promote proper protein folding (3). These enzymes are all large multisubunit complexes (1–3), and they serve not just to catalyze the relevant chemical reactions, but are also critical in regulating protein levels and function. Molecular machines are involved in a host of other processes: for instance, RuBisCo is critical for photosynthesis in algae and plants (4), ATP Synthase is crucial for energy production in cells (5) and the microtubule is critical for cellular infrastructure and organization (6). It is clear that the proper functioning of these machines is critical to essentially all cellular life. Interestingly, however, cells cannot synthesize these machines as functioning units. Instead, cells synthesize the molecular parts that must then assemble in the functional multisubunit complex. Understanding how these machines assemble is thus crucial to understanding how these machines function and how they are regulated.

Since these complexes often perform central roles in cell function and metabolism, they also tend to be associated with a variety of diseases and conditions. For instance, dysregulation of proteasome function has been associated with diseases ranging from cancer to Alzheimer’s (7–11). As a result, understanding the assembly of these complex machines can provide not only a deeper understanding of cell function, but also insights into disease processes. Finally, over the past 10 years, approaches have emerged that allow for the design of synthetic protein complexes that could prove useful in applications as diverse as targeted drug delivery and gene therapy (12–14). Studying the assembly of natural complexes can provide insights into general “design principles” that could allow for the design of synthetic machines that self-assemble more efficiently. Improving our understanding of macromolecular assembly thus has applications ranging from basic cell biology to applied synthetic biology. Advancing our understanding of macromolecular assembly thus holds implications spanning from fundamental cell biology to practical synthetic biology applications.

A common structural motif observed in many macromolecular machines are either rings or contain rings (15–17). Some structures are simple rings: glutamine synthetase (18, 19), pyruvate kinase (20), the KaiC circadian clock protein (21), and AAA proteins (22, 23) are all examples. Many complexes, however, involve rings that are stacked on top of one another. The prototypical example of this type of structure is the proteasome Core Particle (CP), which consists of four rings stacked on top of each other; bacterial proteases like ClpXP/ClpAP and HslUV share a similar architecture (24). The GroEL/ES chaperonin complex and enzymes like the nicotinamice mononucleotide adenylyltransferase and some E2 ubquitin conjugating enzymes also demonstrate a stacked ring architecture (25). In synthetic designs, some complexes are ring-like such as viral capsids (26–29) and other nanorings (30). Rings and stacked rings are clearly a common motif in the evolution of macromolecular machines (16, 17).

Experimentalists have studied the assembly of some machines, particularly the proteasome (2, 31, 32), the ribosome (33, 34) and the GroEL/ES chaperonin (35–37). These studies have resulted in proposed assembly pathways that provide helpful pictures that summarize experimental data. While these pathways are generally well-accepted in their respective communities, they have several limitations—perhaps most notably, they cannot make detailed, quantitative predictions regarding assembly dynamics. It is also difficult to understand, in many cases, why certain aspects of these pathways have evolved, or how particular pathways influence assembly yields. As such, mathematical and biophysical models can be helpful in augmenting these experimental investigations. To date, the vast majority of computational models of self-assembly have focused on the assembly of viral capsids at various levels of biophysical resolution (38–45). While some work has been done using molecular dynamics to study the assembly of molecular machines like the ribosome (46–50), there has been comparatively little work investigating the assembly of rings and stacked rings like the proteasome. One difficulty is time scale; under standard *in vitro* assembly conditions, the proteasome CP self-assembly reaction takes about 3 hours to reach completion (31, 51), which is well beyond the reach of currently-available biophysical simulation techniques (52, 53). It is thus critical to develop computational and mathematical models of assembly that allow us to systematically explore assembly dynamics on realistic timescales.

A little over ten years ago we introduced a mathematical model of the assembly of ring-like structures that attempted to overcome some of the limitations mentioned above (15). In that work we used chemical reaction network theory to develop Ordinary Differential Equation (ODE)-based models of assembly dynamics. Using these models, we showed that ring-like structures can suffer from a severe form of kinetic trapping that we termed *deadlock*, which occurs due to the formation of incompatible stable intermediates. Specifically, when the binding affinities between the subunits is very strong, the subunits quickly form large intermediate structures that cannot react with one another due to steric clashes. These intermediates consume all the monomers, leading to an impasse where no further assembly can occur until the system reaches timescales where these intermediates start to dissociate. The stronger the interactions between the subunits, the longer it takes to for this deadlocked state to resolve, which suggested an evolutionary pressure to avoid very strong interactions in these types of structures. Interestingly, this model predicted that *heteromeric* rings would tend to have a single interaction that was weaker than the other two, and analysis of available experimental structures found that this is indeed the case for most heteromeric rings (15). While this model was highly informative, it was limited to simple ring-like structures.

In this work, we extend this previous model to the important case of stacked ring structures, which, as described above, are an extremely common motif in macromolecular machines (e.g. the proteasome CP, GroEL, etc.). Our main finding is that deadlock in these cases can be significantly worse than in the previous case of single rings—while deadlock in a single ring might typically resolve on timescales of minutes to hours, deadlock for stacked rings can last for days or even longer. We showed that deadlock is worst when all of the interfaces in the structure have high binding affinities. This is true not only in models where there is a fixed initial concentration of subunits (modeling an “*in vitro*” assembly reaction (31, 50, 51) but also in more realistic “*in vivo*” models where we include constant synthesis of subunits or degradation of all intermediates formed (15). These results led us to predict that evolutionary pressures would select against stacked trimers having strong binding affinities both within and between rings. We tested our prediction by analyzing solved stacked trimer structures; we found that indeed the majority – if not all – of the stacked trimers did not contain two very strong interactions. In experimental studies of assembly, it is common for researchers to discuss the notion of “assembly pathways,” but to our knowledge there has been no attempt to rigorously define what an assembly pathway actually is. In order to better understand the origins of deadlock and why certain patterns of interaction affinity generate a deadlocked state, we defined an assembly pathway as a binary tree representing a scenario of how a group of subunits go from being monomers to form the fully assembled structure. Our assembly pathway analysis showed that when all the binding affinities are strong, many of the possible pathways are utilized during assembly, consuming subunits and creating high levels of deadlock. In contrast, when one of the interfaces is weaker, only a small number of pathways contribute significantly to assembly, suggesting that observed structures have evolved patterns of interaction affinities that enforce a more hierarchical assembly process (15). In sum, our work provides critical insight into the evolutionary pressures that have shaped the assembly of stacked rings. Furthermore, our work posits a potential design principle for to optimize self-assembly efficiency in stacked rings: synthetic structures should likely be designed not to have strong interactions throughout but instead have a mixture of binding strengths to avoid deadlock during assembly. Since previous design approaches typically attempt to maximize the affinity of every interface (54–58), adopting a different approach may lead to a higher success rate for designing such structures. Thus this work highlights the role of mathematical modeling in understanding self-assembly processes in evolution, biomedicine, and synthetic biology.

## Results

### Constructing a model of stacked ring assembly

In this work we model the self-assembly of a stacked ring-like protein complex containing *n* subunits like the one depicted in **Figure 1A (left)**. As an example, in the X-ray structure from Protein Data Bank (PDB ID 2FO3) in **Figure 1A (left)** each subunit is depicted with a unique color. We consider a stacked trimer containing two 3-membered homomeric rings (*n* = 6). To construct a model of a stacked trimer, we can visualize each subunit a node in a graph and a stacked ring as two connected symmetric trimeric rings (**Figure 1A, right**), where the connections are representative of the noncovalent bonds between subunits. Stacked trimers have a 3-fold axis of rotational symmetry as well as mirror symmetry between rings, so that the bottom ring is the mirror image of the top ring (16, 17). Indeed, there are examples of homomeric stacked rings such as Nicotinamide mononucleotide adenylyltransferase (59), Periplasmic Heme-binding Protein ShuT (60), and Glycerophoshphodiesterase (61) to name a few.

**Figure 1:**
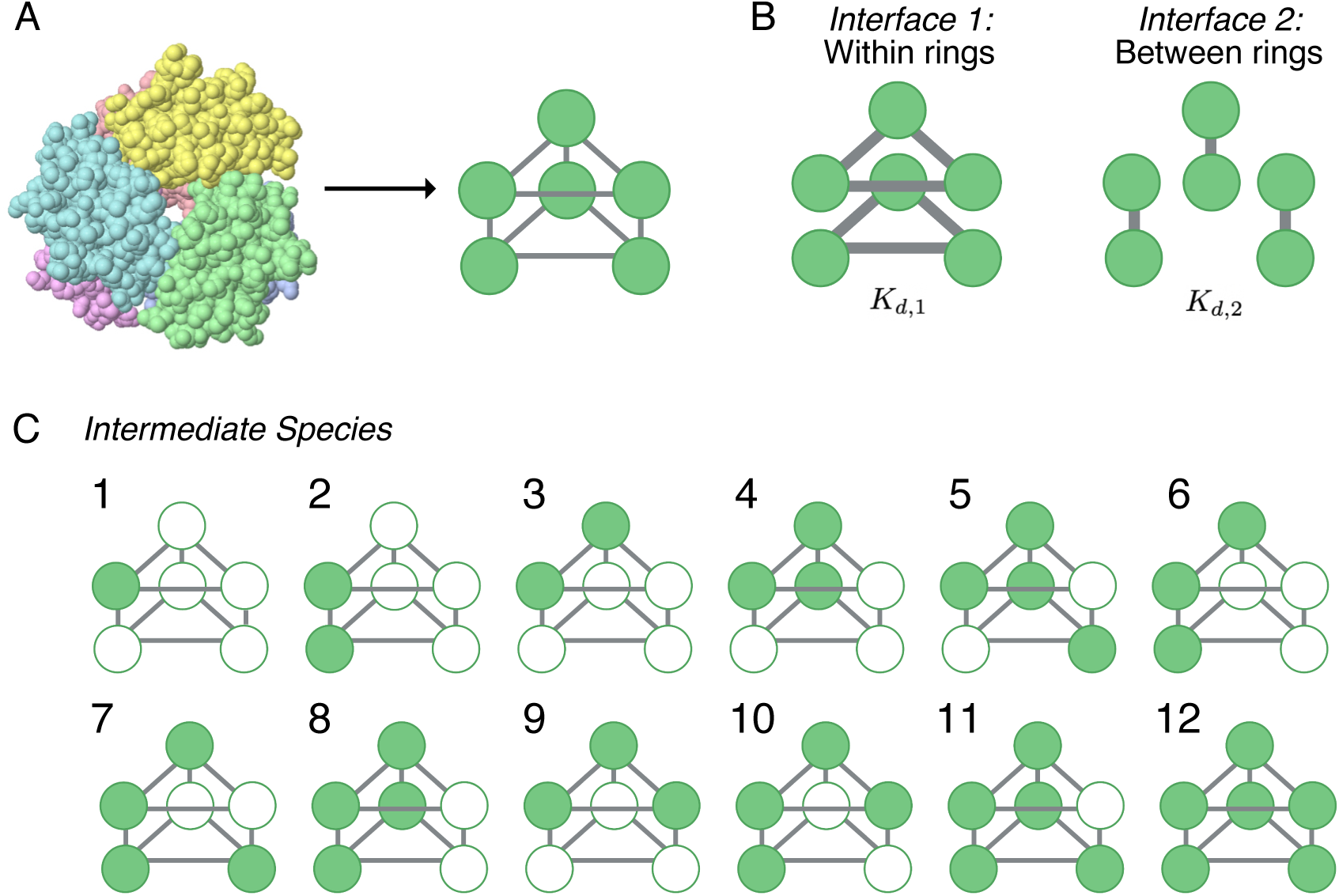
Stacked trimer model schematic. **(A)** An example stacked trimer (X-ray structure from Protein Data Bank, PDB ID 2FO3) on the left is depicted with each monomer in a unique color. Any such stacked trimer can be represented as a graph with edges representing the non-covalent bonds between the 6 proteins. Each protein has three binding interfaces (bonds are represented by the solid gray lines). **(B)** Each protein can form two types of bonds, with proteins on the same ring (interface 1: within) and with proteins from the opposing ring (interface 2: between). Each bond type has associated *K*_*d*_ values, *K*_*d*,1_ and *K*_*d*,2_, respectively. **(C)** A list of all the intermediate species that can be formed during stacked trimer assembly in our model.

Similar to a previous model of single ring assembly (15), the subunits are treated as identical but have a “sidedness,” that is, each subunit has three distinct interfaces: left, right, and across. The left side of one subunit can bind to the right side of another and we call these the *within-ring interactions* (**Figure 1B, left**). The across side of one subunit can bind to the reflecting across side of another, in an up-down fashion, giving rise to what we call *between-ring interactions* (**Figure 1B, right**). Thus, each monomer contains three types of binding interfaces: two for binding within the ring (the left and right side of what we call *interface* 1) and one for binding between the rings (which we call *interface 2*, **Figure 1B**). For each bond type, there is an associated binding affinity. For *interface 1*, the binding affinity between each subunit is described by a dissociation constant *K*_*d*,1_ and for *interface 2*, the binding affinity between the two subunits forming the between-ring bond is *K*_*d*,2_. Note that these *K*_*d*_ values are defined based on the association and dissociation rates of the chemical reactions that form the corresponding dimers from the monomers (see **Supplemental information** for further details).

We call any structure that is smaller than the stacked trimer an “intermediate,” that is, all intermediates are sub-structures of the fully assembled stacked ring. Formally, an intermediate is any connected subgraph of the fully assembled structure. As such, intermediates have a number of subunits between 1 and *n* − 1. We have enumerated all the possible intermediate structures for the stacked trimer in **Figure 1C**. Since every intermediate is a substructure of the fully-assembled complex, we use a simple visual language for these intermediates. The filled-in circles represent subunits that are “present” in that structure, whereas unfilled circles represent subunits that are “absent” from that intermediate. Note that the unfilled circles represent all the available locations in the intermediate where other intermediates can bind. To enumerate the species, we first start with Species 1, which is simply the monomer. Then, we add another monomer in every possible available location on the graph and determine whether the resulting structure forms a connected component. If it does, this is a new intermediate and we add it to the list. Doing so generates the two possible dimer species, Species 2 and 3. Iterating this process of adding monomers to the two dimers generates all possible trimers, and continuing this process allowed us to generate all 11 possible intermediates in **Figure 1C**. We denote an individual species “*i*” as *S*_*i*_.

To construct a mathematical model of stacked trimer assembly, we next enumerated all the binding reactions between intermediates. To do so, we checked all possible pairwise combinations of the intermediates in all possible relative orientations (see **Supplemental Material Section 1**) to see if they could react to form another intermediate or the fully-assembled structure. For example, consider Species 1 and 2 (*S*_1_ and *S*_2_), the monomer and the “*K*_*d*,2_” dimer. If we rotate Species *S*_1_ by 120°, note that it “fits into” an open spot in the Species *S*_2_ structure. As a result, there is a reaction between *S*_1_ and *S*_2_ that generates another intermediate, in this case *S*_6_. Similarly, if we rotate *S*_1_ by 240°, that induces a reaction that generates *S*_4_ (See **Supplemental Figure S1**). Using this basic approach, we enumerated all the possible binding reactions that could occur between intermediates. The complete list may be found in **Supplemental Material Section 1.2-1.4**.

The concentration of each species is represented by *X*_*i*_ where *i* is the index of the intermediate and not the number of subunits in the structure. Note that this value is in general a function of time (i.e. *X*_*i*_(*t*)); in our notation, however, we leave the dependence on time implicit. The number of subunits in an intermediate is given by *η*(*S*_*i*_), which is a function that maps from the species index to the number of subunits present in the given species. For example, *S*_1_ is the monomer (**Figure 1C**) and *η*(*S*_1_) = 1 since there is only one subunit present. In contrast, *S*_10_ has *η*(*S*_10_) = 4 since there are 4 subunits present in that intermediate. Additionally, we define 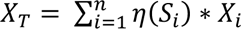 to be the total concentration of the subunits (i.e. the total concentration of all subunits in all the species combined).

Similar to previous models (15, 62, 63), we used the standard law of mass action to derive a system of Ordinary Differential Equations (ODE’s) that describes the time evolution of the concentration of all the species in the model based on these reactions (**Figure 1C**). The derivation is described in detail in **Supplemental Material Section 1.5**. In this work we make the simplifying assumption that the association rate between intermediates is the same, regardless of the size and structure of the intermediates in question. All association reactions in our model thus occur at the same rate, which we term *k*_+_. All binding reactions in the model are of course reversible, and the dissociation rates depend on both the number and the strength of the noncovalent bonds being formed between and/or within each ring. As in previous models of self-assembly, we assume the intermediate structures are fairly rigid (15, 64). This allows us to use straightforward thermodynamic arguments to calculate the dissociation constant for any given reaction, which we call *K*_*eff*_. This calculation is described in detail in the **Supplemental Material Section 1.4** and is based on the number of each type of interface that is disrupted for a given dissociation reaction. Note that, as described previously, ring-like structures are very stable, and so dissociation reactions that disrupt more than one noncovalent bond happen at very low rates (15, 64, 65). Using these assumptions, and straightforward calculations of how many “ways” two intermediates can combine to form another (See **Supplemental Material Section 1.5**), we derived a system of ODEs for the self-assembly of the stacked trimer. These ODE’s were numerically solved using Python 3.9.7 for fixed parameter values (See Methods). All of the code used for this work is available at: https://github.com/llagunes-324B21/stackedTrimer_detModel.

We define the *assembly yield* of an intermediate *i* as 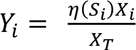 for each species *i* = 1, 2, …, 12 at a given time point *t*. Note that *Y*_*i*_ is simply the fraction of total monomers in the system that are found in that given intermediate. Take, for instance, *Y*_12_, the fraction of monomers in the fully-assembled stacked trimer. If *Y*_12_ = 1 at some point in time, that would mean that 100% of the monomers are found in the stacked trimer structure; if *Y*_12_ = 0.5, then only 50% of the monomers are found in the fully-assembled structure. When *Y*_12_ < 1, this indicates that some monomers are sequestered in smaller intermediates. We use this definition of the assembly yield to investigate the role of the binding affinities on the formation of the stacked trimer.

### The assembly dynamics of stacked homomeric rings

To investigate the role the binding affinities *K*_*d*,1_ and *K*_*d*,2_ have on the formation of the stacked trimer assembly yield, we simulated the dynamics obtained from our model of stacked ring assembly starting from an initial condition of only monomers. As in previous work, in order to parameterize our model, we made the simplifying assumption that the rates of all the association reactions (i.e. the rate constants for all of the binding reactions) are the same, regardless of the identity of the reactants (15). We have taken the association rate to be *k*_+_ = 10^6^ M^-1^ s^-1^, which is a reasonable value for protein-protein interactions (15, 66). As described previously, changing the value of this rate-constant in the model corresponds to a simple re-scaling of the time units of the simulation (15). Thus, while the exact timescale depends in the specific value of the association rate, the general shape of the curves discussed in this work do not depend on the value of *k*_+_. For our first exploration of self-assembly dynamics in this model, we used an initial monomer concentration of 4 μM, i.e. *X*_1_ = 4 × 10^-6^ M.

We first consider the scenario for which the total concentration of subunits *X*_*T*_ is fixed. In other words, there is no synthesis of new monomers, or any intermediate, and there is no protein degradation. The specific system of ODEs corresponding to this scenario may be found in **Supplemental Material Section 1.5**. We call this case the *in vitro* model to resemble *in vitro* biochemical experiments where self-assembly occurs in a solution has fixed total protein concentration (15, 31, 32). **Figure 2A** shows the assembly yield dynamics of the stacked trimer for a set of unique combinations of *K*_*d*,1_ and *K*_*d*,2_. In this model, we find that the assembly dynamics depend critically on the affinity between the subunits. When both affinities are extremely strong (*K*_*d*,1_ = *K*_*d*,2_ = 10^-12^ M), the overall yield is relatively low, reaching at most 0.5 (i.e. 50% full assembly) even on relatively long timescales (10^6^s, or around 11 days) (**Figure 2A**, green curve). As mentioned in the introduction, the fundamental reason for this is a type of kinetic trapping that we have termed “deadlock” (15). In this parameter regime, the monomers react very quickly with one another to form various larger intermediates, and, because of the strength of the interactions between the subunits, those intermediates are relatively stable. Most of them also cannot react with one another due to steric clashes. For instance, when *K*_*d*,1_ = *K*_*d*,2_ = 10^-12^ M, the system is dominated by mostly Species 11 and Species 7 during the “plateau” phase of the dynamics (**Supplemental Material Section 2, Figure S6**). These intermediates cannot react with one another; for example, Species 8 (**Figure 1B**) cannot react with any other structure but a dimer or a monomer. Since the monomers and dimers are rapidly depleted in this parameter regime, Species 8 cannot react with anything, thus leading to deadlock.

**Figure 2:**
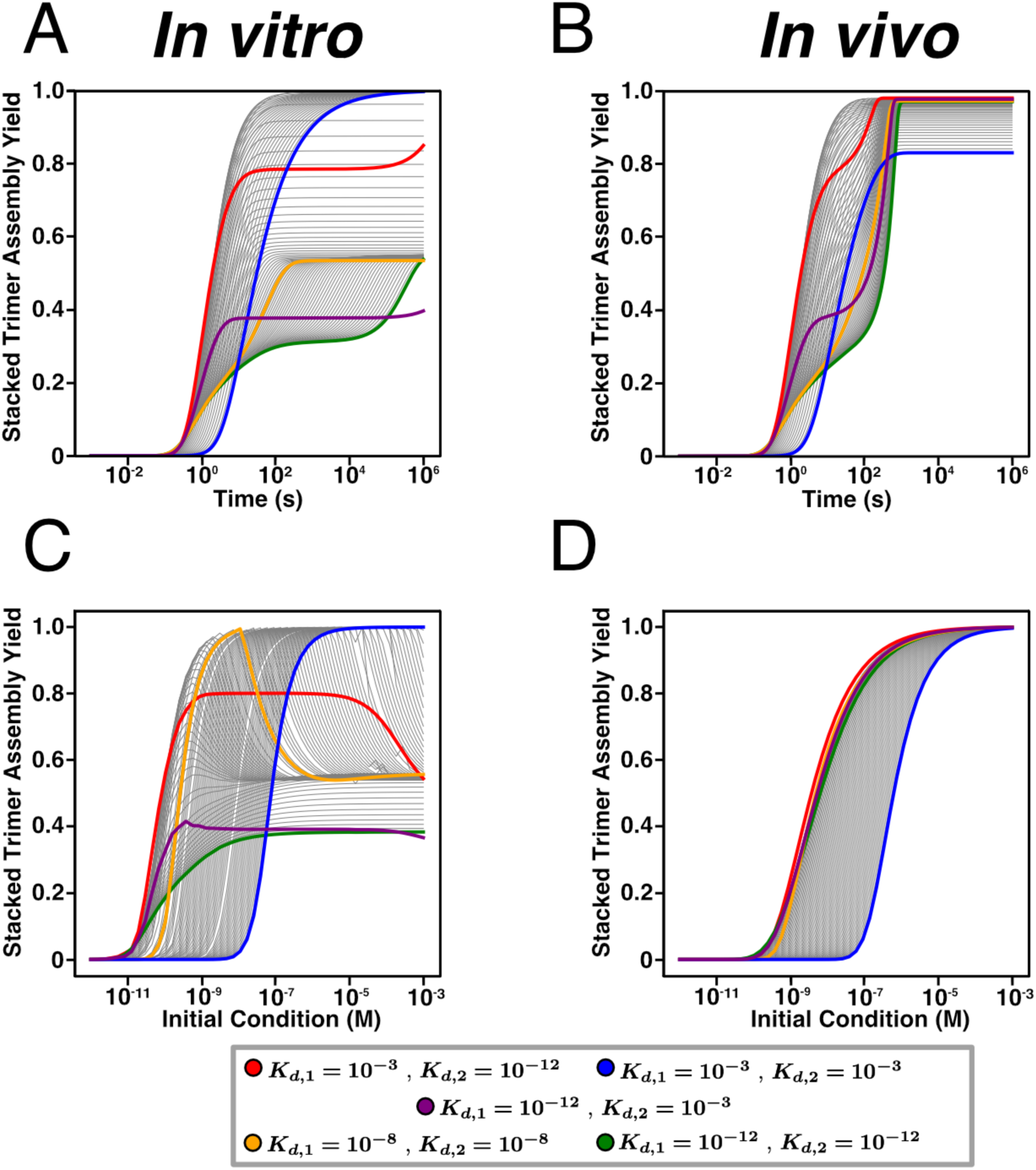
Dynamics of stacked trimer assembly. **(A)** Time course plot for *in vitro* and **(B)** *in vivo* model with different *K*_*d*,1_ and *K*_*d*,2_ parameter values. Initial monomer concentration is 4 × 10^-6^ M, and for the *in vivo* model the degradation rate δ is set to 2.8 × 10^-4^ s^-1^. **(C-D)** Assembly yield curves with respect to increasing initial monomer concentration with different *K*_*d*,1_ and *K*_*d*,2_ parameter values. Same color scheme as in **(A-B)** for both *in vitro* and *in vivo* models.

In order for deadlock to resolve, larger intermediates must dissociate to generate smaller species that can then react (15). In the case of the stacked trimer, many of the intermediates are actually rings themselves (e.g. Species 8 and 9). For simple thermodynamic reasons, rings are far more stable than chains (15, 64), and so this dissociation timescale can be theoretically quite long. As a result, while deadlock resolves in reasonable timescales for the case of the assembly of simple rings, it can last well beyond biologically reasonable timescales in the case of the stacked trimer (Supporting Information) (**Figure 2A**).

In **Figure 2A**, we have highlighted several other examples of combinations of binding affinities that give rise to interesting and characteristic assembly dynamics. When both affinities are of intermediate strength (*K*_*d*,1_ = *K*_*d*,2_ = 10^-8^ M, **Figure 2A**, yellow curve), we see similar levels of deadlock to the case of two extremely strong interactions discussed above. Making one of the interactions much weaker produces strikingly different results depending on which interaction is strong and which is weak. If the interaction *within* the rings is strong (*K*_*d*,1_ = 10^-12^ M, *K*_*d*,2_ = 10^-3^ M, **Figure 2A**, purple curve), then we see even more extreme deadlock than when both interactions are strong. This is because the species with the highest concentration during the deadlock phase is Species 11, the intermediate with only one monomer missing. Additionally, there is a relatively high concentration of Species 9 which is the single ring structure. These two structures are not compatible and therefore cannot form a stacked trimer, which lead to deadlock and since the binding affinity *within* the rings is very high, then Species 9 is not disassembling quickly enough to overcome deadlock (**Supplemental Material Section 2, Figure S6)**. In contrast, if the *between-ring* interaction is strong (*K*_*d*,1_ = 10^-3^ M, *K*_*d*,2_ = 10^-12^ M, **Figure 2A**, red curve), the impact of deadlock is much less. This indicates that it is not the total strength of the interactions within the structure, but rather the *pattern* of interaction strengths, that dictates the dynamics of assembly in stacked rings. Finally, when both interactions are very weak (*K*_*d*,1_ = *K*_*d*,2_ = 10^-3^ M, **Figure 2A**, blue curve), the system completely avoids deadlock, but the assembly dynamics are quite slow. That is because the smaller intermediates, like the two dimer species, are much less stable and thus it takes considerably longer for larger intermediates to form (**Supplemental Material Section 2, Figure S6)**.

The above model considers the case of an *in vitro* biochemical experiment, and it may be applicable to situations inside cells where monomeric species are pre-formed and then assemble in response to some incoming signal (67–70). In the case of many molecular machines, however, we anticipate that monomers would be constantly synthesized, and assembly takes place to replace structures that are lost due to degradation or dilution from cell division (71). We model this by introducing a constant synthesis rate of monomers and a first-order degradation term for all the intermediates; we call this the “*in vivo*” model since. We fixed the degradation term *δ* to be 2.8 × 10^-4^s, which models the case where, say, a bacterial cell divides approximately once very hour (15). In our *in vivo* models we did not consider cases where the total protein concentration changes over time. We thus set the monomer synthesis rate *Q* = *δ* · *X*_*T*_ (**Supplemental Material Section 1.5**). **Figure 2B** shows the assembly yield trajectories for the stacked trimer for combinations of *K*_*d*,1_ and *K*_*d*,2_. The curves represent the same *K*_*d*,1_ − *K*_*d*,2_ parameter combinations as in **Figure 2A** with the same color scheme. In this scenario, the assembly yield of the stacked trimer is relatively high for most *K*_*d*,1_ and *K*_*d*,2_ values. Deadlock is also less severe than seen in the *in vitro* model—while there are “plateaus” in the assembly dynamics (Figure 2A, green, purple and red curves), these plateaus resolve on much faster timescales than in the *in vitro* case. This is because monomers are being constantly synthesized, and thus the system never truly “runs out” of them as it does in the *in vitro* model. The primary impact of deadlock in this case is the fact that the steady-state yield depends strongly on the pattern and strength of binding affinities. For instance, weak interactions lead to relatively low yields (**Figure 2B**, blue curve). Interestingly, having uniformly strong interactions (*K*_*d*,1_ = *K*_*d*,2_ = 10^-12^ M) leads to lower steady-state yields than cases where one of the interactions is weak (e.g. *K*_*d*,1_ = 10^-12^ M, *K*_*d*,2_ = 10^-3^ M, **Figure 2B**, comparing the green and purple curves). This suggests that, as in the case of single rings, deadlock can impact steady-state yields even in models where monomers are constantly synthesized in the cell (15).

In the models thus far, the initial monomer concentration is fixed at 4 μM, i.e. *X*_1_ = 4 × 10^-6^ M. To characterize how the assembly yield of the stacked trimer responds to changes in the initial monomer concentration, we first considered the *in vitro* model, where there is no protein synthesis or degradation. In **Figure 2C**, we calculated the assembly yield of the stacked trimer after 24 hrs for different initial monomer concentrations and fixed *K*_*d*,1_ and *K*_*d*,2_ values. Note that in **Figure 2C**, the colored curves correspond to the same *K*_*d*,1_ and *K*_*d*,2_ combinations as in the previous panels. When both affinities are of intermediate strength (*K*_*d*,1_ = *K*_*d*,2_ = 10^-8^ M, **Figure 2C**, yellow curve), we see that the assembly yield for the stacked trimer increases as the initial monomer concentration increases, reaches a peak, and then begins to decrease with increasing initial monomer concentration. In the first phase before the peak, the initial monomer concentration is low and thus there is slow binding between the intermediate structures. As a result, it takes longer for the stacked trimer to be assembled, leading to low assembly yields after 24 hrs. In the second phase after the peak, the initial concentration is high and yet the assembly yield for the stacked trimer decreases. This is because there is now fast binding between the intermediates due to the higher protein concentration. The rapid binding of monomers and other smaller intermediates leads to deadlock, as described above (**Figure 2A**). At the peak, a balance is achieved for this particular pattern of interaction affinities: binding is not too fast, and so the system is able to avoid deadlock and achieve near-100% yield.

As with the assembly kinetics described in **Figure 2A**, the response of the system to changes in initial conditions depends critically on the pattern of binding affinity within the structure. For instance, when both interactions are very strong (*K*_*d*,1_ = 10^-12^M, *K*_*d*,2_ = 10^-12^M, **Figure 2C**, green curve), we see an increase in assembly yield up to about 40%. The interactions are so strong in this case that the system essentially cannot avoid deadlock no matter the initial concentration of monomers. If we make the *between-ring* interaction much weaker (*K*_*d*,1_ = 10^-12^M, *K*_*d*,2_ = 10^-3^M, **Figure 2C**, purple curve) we see essentially the same behavior, indicating that weakening the bonds between the rings does not make the system more robust to deadlock. Interestingly, however, if we reverse the situation so that the *between-ring* interaction is strong and the *within-ring* interaction is weak (*K*_*d*,1_ = 10^-3^ M, *K*_*d*,2_ = 10^-12^ M, **Figure 2C**, red curve), we see the assembly yield for the stacked trimer increase to a maximum level and then decrease with increasing initial monomer concentration. In this case, the maximum assembly yield for the stacked trimer spans a longer region of initial monomer concentrations than in the intermediate strength (*K*_*d*,1_ = *K*_*d*,2_ = 10^-8^ M) case. However, that optimal region reaches approximately 80% stacked trimer assembly, suggesting that the system cannot fully avoid deadlock. Finally, when both affinities are weak (*K*_*d*,1_ = 10^-3^ M, *K*_*d*,2_ = 10^-3^ M, **Figure 2C**, blue curve), the system reaches 100% assembly at 24 hrs, but only when protein concentrations are relatively high. This is due to the fact that, while binding occurs quickly at higher monomer concentrations, the intermediates formed are unstable (particularly Species 2-6, **Figure 1C**) and dissociate before further assembly reactions can occur. Under these conditions, assembly is very slow and higher protein concentrations are required to achieve 100% yield. Note that, even when both bonds are very weak, at even higher protein concentrations yields begin to decrease due to deadlock (**Supplemental Material Section 1.5**). Non-monotone yield curves as a function of protein concentration are thus characteristic of *in vitro* assembly of the stacked trimer, which is different from the behavior of single rings (15).

Changing total protein concentration in the *in vivo* model is not as straightforward as the *in vitro* case. As discussed above, the total protein concentration in this model is just *X*_*T*_ = *Q*/*δ*, i.e. the ratio of the monomer synthesis rate to the first-order degradation rate for all the species. One can thus change the total protein concentration by changing either the value of *Q* or *δ*. In this case we chose to modify *Q*, since this is most closely analogous to increasing total monomer concentration as in **Figure 2C**. This also models a case where, for instance, a bacterial population is growing at a constant rate (in this case, doubling every hour) and is attempting to increase the concentration of the stacked monomer by increasing the synthesis of the monomers. In **Figure 2D**, we show the steady-state yield of stacked trimers as a function of total protein concentration, highlighting particular patterns of interaction affinities with distinct colors as with all the other panels in the figure. In this case we do not observe non-monotone behavior—the higher the value of the monomer synthesis rate, the higher the steady-state yield. The pattern of affinities within the structure, however, still has a large impact on assembly. When both interactions are very weak (*K*_*d*,1_ = *K*_*d*,2_ = 10^-3^ M, **Figure 2D**, blue curve), assembly yield does not reach 100% until relatively high monomer concentrations. Interestingly, however, having both interactions very strong (*K*_*d*,1_ = *K*_*d*,2_ = 10^-12^ M, **Figure 2D**, green curve) generally results in lower assembly yields at any given total monomer concentrations than the other scenarios, particularly the cases where both interactions are of intermediate strength or the *between-ring* interaction is strong (**Figure 2D**, red and yellow curves).

The results in **Figure 2D** suggest that there may be strong evolutionary pressures that impact the patterns of affinities in stacked ring structures. Increasing the value of *Q* represents a direct increase in the amount of *energy investment* the cell is making in the synthesis of monomers. The higher this value is, the higher the total protein concentration is, and the more ATP the cell is spending on the synthesis of these proteins. If we assume that only the fully-assembled structure is functional, as is the case for the proteasome, GroEL (37, 72), and a host of other structures (73–78) then the assembly yield represents the percentage of that energy investment that actually results in functional machines that can support cell physiology. In other words, this is the return on that energy investment. We would expect that there would be some evolutionary pressure to maximize that return in many cases, particularly in bacterial species where even a small waste of energy can have detectable fitness consequences (79, 80). Since some curves are always near the top of **Figure 2D**, that suggests certain patterns of affinity almost always result in higher yields, regardless of the total protein concentration present, and thus would likely be evolutionarily favored.

To further assess the effect of the binding affinities on yield for the stacked trimer, we generated heat maps for both the *in vitro* and *in vivo* models. **Figure 3A** shows the assembly yield for the *in vitro* model after 24 hours for values of *K*_*d*,1_ and *K*_*d*,2_ ranging from 10^-3^*M* to 10^-12^*M* and a fixed initial concentration at 4 μM. The contour lines show highlighted assembly yield values. This type of plot allows us to visualize the effect of the binding affinities on assembly yield. If both *K*_*d*,1_ and *K*_*d*,2_ are weak (upper right quadrant of the plot), we see that the assembly yield is greater than 95%. Interestingly, the lowest assembly yield occurs in the black-colored region delineated by the 40% contour line; this occurs when both *K*_*d*,1_ and *K*_*d*,2_ are both strong (lower left quadrant) and when the *between-ring* interaction is much weaker than the *within-ring* interaction (i.e. when *K*_*d*,1_ is strong and *K*_*d*,2_ is weak). These results further highlight that assembly is vastly impacted by deadlock in the *in vitro* model when both interactions are strong, or when *K*_*d*,1_ is strong. **Figure 3B** shows a similar heat map for the steady-state yield of the stacked trimer in the *in vivo* model. In this case, the degradation rate remains fixed at *δ* = 2.8 × 10^-4^ s^-1^, the initial monomer concentration is also 4 μM and we ranged *K*_*d*,1_ and *K*_*d*,2_ from 10^-3^*M* to 10^-12^*M*. We see that when both *K*_*d*,1_ and *K*_*d*,2_ are strong (lower left region), the assembly yield for the stacked trimer is not optimal. However, the lowest assembly yield occurs when both *K*_*d*,1_ and *K*_*d*,2_ are weak (upper right region). Note that overall, the assembly yield is higher for all values of *K*_*d*,1_ and *K*_*d*,2_ in comparison with the *in vitro* model with the lowest assembly yield at approximately 82% in the *in vivo* model and 30% in the *in vitro* model, similar to the results in **Figure 2A** and **Figure 2B**. Since both models show that when both binding affinities are strong the assembly yield for the stacked trimer is non-optimal, this suggests that there is an evolutionary pressure to select other combinations of binding strengths. In other words, we predict that stacked trimer structures would not tend to have strong binding affinities both *within* and *between* the rings.

**Figure 3:**
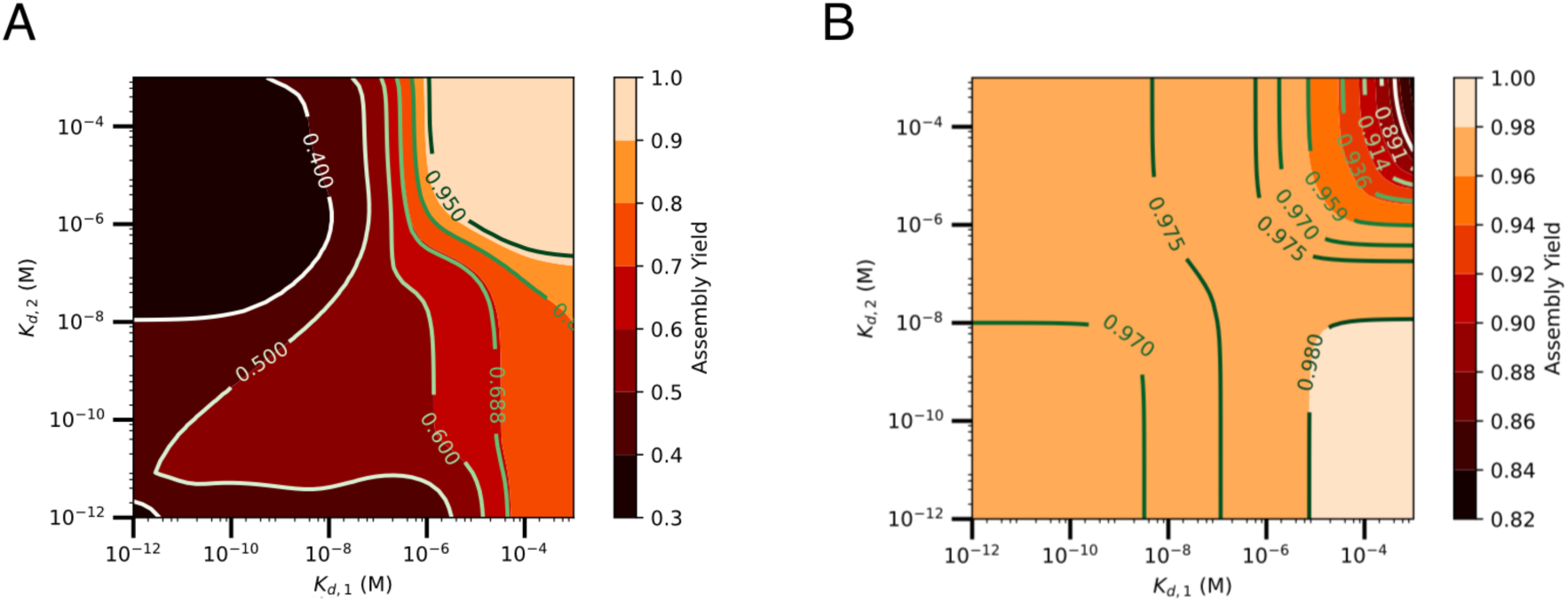
Assembly yield for different values of *K*_*d*,1_ and *K*_*d*,2_. **(A)** Heatmap showing assembly yield after 24 hours for *in vitro* and **(B)** *in vivo* models with fixed initial monomer concentration of 4 × 10^-6^ M and the binding affinities *K*_*d*,1_ and *K*_*d*,2_ increase from 10^-12^ to 10^-3^. Note that all results for the *in vivo* case correspond to steady-state yields. Contour lines are shown to visualize assembly yield values. For the *in vivo* model, *δ* = 2.8 × 10^-4^ s^-1^.

### Observed stacked trimer structures avoid having two strong interactions

The findings above suggest that there is likely an evolutionary pressure for stacked rings to not exhibit strong binding affinities both *within* and *between* rings. To test this prediction, we considered solved structures of stacked trimeric rings. We used the PDBePISA database, which contains structural and chemical properties of macromolecular surfaces, interfaces and assemblies for all structures in the PDB. We began by assembling a set of over 3000 homo-hexameric structures. Since PDBePISA does not annotate the general geometry of structures, we visually inspected each of these complexes to collect only those that form stacked trimeric rings. In order to avoid misclassifications, we defined a stacked trimer as one that has a similar topology to the one in **Figure 1A (right)**, where there are three bound monomers on a top plane bound to three bound monomers on a bottom plane. Additionally, a stacked trimer would need to have rotational symmetry along each ring and mirror symmetry between the rings (see **Supplemental Material Section 3**). In other words, we removed any structures that were single rings, like those discussed in (15), chains or structures that did not match the topology considered in our model. This resulted in 1580 stacked ring structures; a list of these structures with their PDB IDs is provided as an additional supplementary file in: https://github.com/llagunes-324B21/stackedTrimer_detModel.

Estimating binding affinities from the structures of protein-protein interfaces is difficult. In our analysis, we focus on the Buried Solvent-Accessible Surface Area (BSASA), which can be used as proxy for binding affinities (15, 81, 82). In this case, if there is little contact between the two, then the BSASA is small and the binding affinity likely weak. Alternatively, if there is a large contact between the two monomers, then the BSASA is a relatively high and the binding affinity is likely stronger. We used the BSASA values calculated in PDBePISA for the interfaces *between* and *within* rings. For any stacked trimer, there are a total of six *within* ring interfaces. For each structure, we used the average of all six, which we call the *K*_*d*,1_ BSASA. Similarly, there are a total of three *between* ring interfaces; the average of these three interfaces is the *K*_*d*,2_ BSASA. The calculated *K*_*d*,1_ and *K*_*d*,2_ BSASA for each structure are shown in **Figure 4A** (each red dot represents one of the 1580 stacked trimer structures). In the scatterplot, we see that the majority of structures tend to cluster in the lower left region where *K*_*d*,1_ and *K*_*d*,2_ BSASA’s are relatively equal and not extremely large, suggesting that the majority of stacked trimers tend avoid very strong interactions. There are some structures that have a low *K*_*d*,1_ BSASA with high *K*_*d*,2_ BSASA. These structures have much more interface contact *between* the rings than *within* the rings. There are also two structures with the opposite pattern, with a high *K*_*d*,1_ BSASA and a low *K*_*d*,2_ BSASA. The fact that structures where the *between* ring interface much stronger than the *within* ring interface are much more common than the alternative matches our prediction that this pattern of assembly affinities leads to higher yields (**Figures 2** and **3**) Interestingly, none of the stacked trimer structures we found in the PDBePISA database show high BSASA for both the *between* and *within* interfaces (green rectangle in **Figure 4A**), consistent with our prediction that two strong bonds would not be selected for in the evolution of stacked trimers.

**Figure 4:**
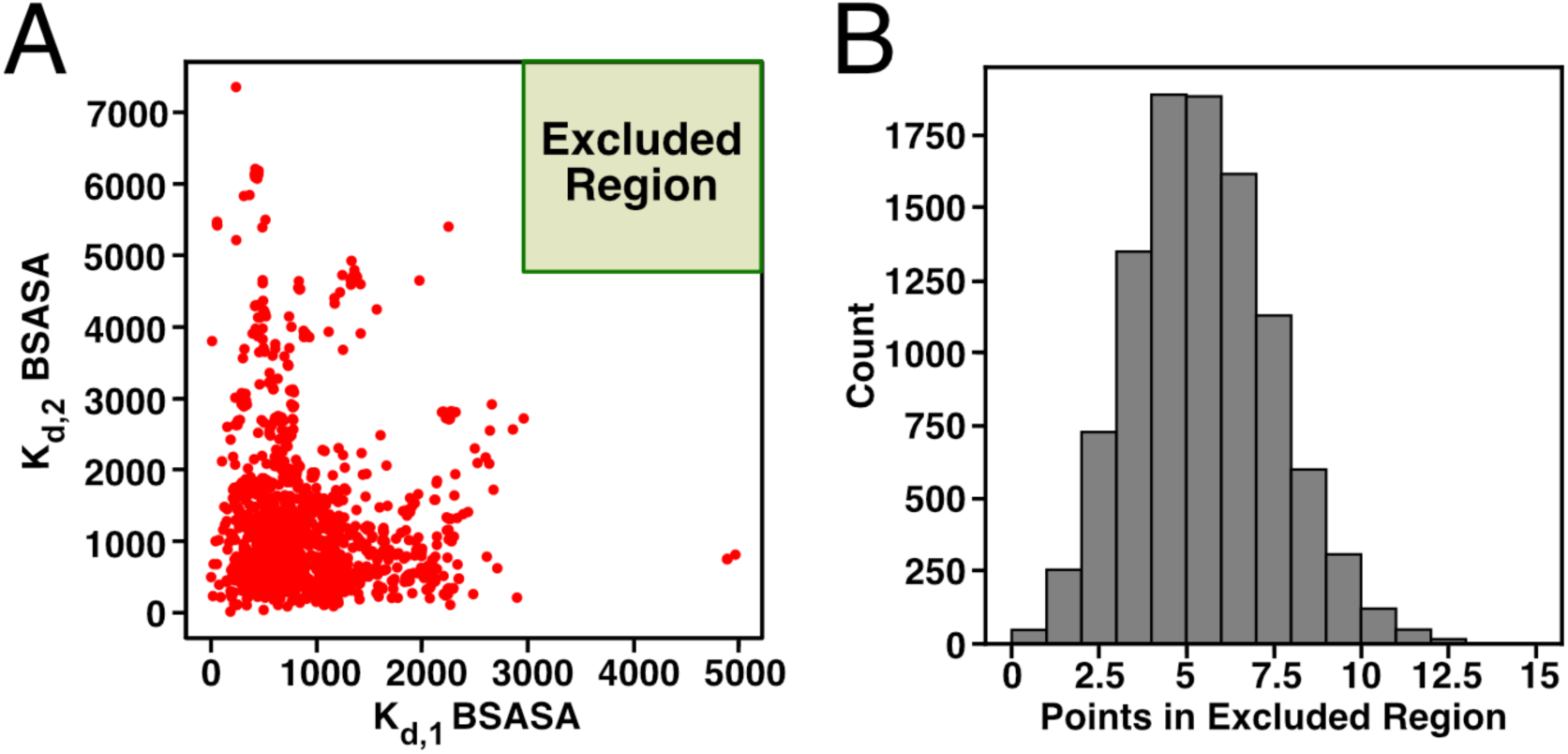
Buried Solvent-Accessible Surface Area (BSASA) in Protein Data Bank stacked trimer structures. (**A**) We identified stacked trimers identified the PDBePISA database; we collected the BSASA for interfaces between and within rings for each. We used the BSASA within and between rings as a proxy for *K*_*d*,1_ and *K*_*d*,2_, respectively. (**B**) From the stacked trimers in (**A**), we resampled the data using a permutation test and determined how many cases had *K*_*d*,1_ and *K*_*d*,2_ BSASA in the excluded region (green rectangle in (**A**)). The histogram shows the total cases where *K*_*d*,1_ and *K*_*d*,2_ BSASA were in the green rectangle (counts) based on 10,000 trials. Note that out of the 10,000 trials, only 48 had 0 cases where *K*_*d*,1_ and *K*_*d*,2_ BSASA were in the excluded region (green rectangle).

Since we did not see any stacked trimers with both high *K*_*d*,1_ and *K*_*d*,2_ BSASA’s, we performed a simple permutation test with 10^4^ replicates to statistically test our prediction that stacked rings do not exhibit strong binding affinities for both *within* and *between* rings. Our permutation test consisted of resampling or “shuffling” the *K*_*d*,1_ and *K*_*d*,2_ BSASA’s between all the structures and checking if any of the shuffled structures were at a threshold of 4000 Å^2^ or higher for both interfaces (green rectangle in **Figure 4A**). In **Figure 4B**, we see the distribution of the number of cases observed in the green rectangle from this test. This distribution shows that only 48 trials had no cases where *K*_*d*,1_ and *K*_*d*,2_ BSASA were in the green rectangle. This suggests that the observation of having no PDB structures with two strong interactions is unlikely to have occurred at random (*p* < 10^-3^). It is thus likely that evolution has avoided this scenario in order to lower the impact of deadlock on both assembly kinetics and steady-state yield.

### Defining and enumerating assembly pathways

Our results thus far show that the assembly of stacked trimers yields deadlock similar to the assembly of single rings (15), and that patterns of binding affinities play a role in that deadlock. It is unclear, however, *why* certain patterns of affinities might lead to higher or lower levels of deadlock. For instance, why would having high affinity for both the *between* and *within* ring interactions yield to lower yields? In order to answer this question, we developed an approach to analyze *assembly pathways*. While the term “assembly pathway” is used broadly in the field (83–87), we are aware of no precise, formal, mathematical definition for what an assembly pathway actually is.

Informally, an assembly pathway should represent a distinct scenario representing how a group of subunits go from being monomers to form the fully assembled structure. Somewhat more formally, we define an assembly pathway to be a strict binary tree where the nodes are assembly intermediates (including the monomers and the fully-assembled structure). An example is shown in **Figure 5A**. In this tree, each node has two *child nodes* and one *parent node*. The exceptions to this are the *leaf nodes,* which have no child nodes, and the *root node*, which has no parent node. The leaf nodes are always monomers, and an assembly pathway has exactly the same number of leaf nodes as there are subunits in the fully-assembled structure (six in the case of the stacked trimer, **Figure 5**). The root node is always the fully-assembled structure. If two nodes are to be child nodes of a given parent, then there must be a chemical reaction between the two child species that generates the parent. In other words, whenever we have two nodes *below* a parent node in the tree, they have to be able to combine to generate that particular intermediate (**Figure 5A**). A more complete and formal definition of assembly pathways is given in the **Supplemental Material Section 4**.

**Figure 5:**
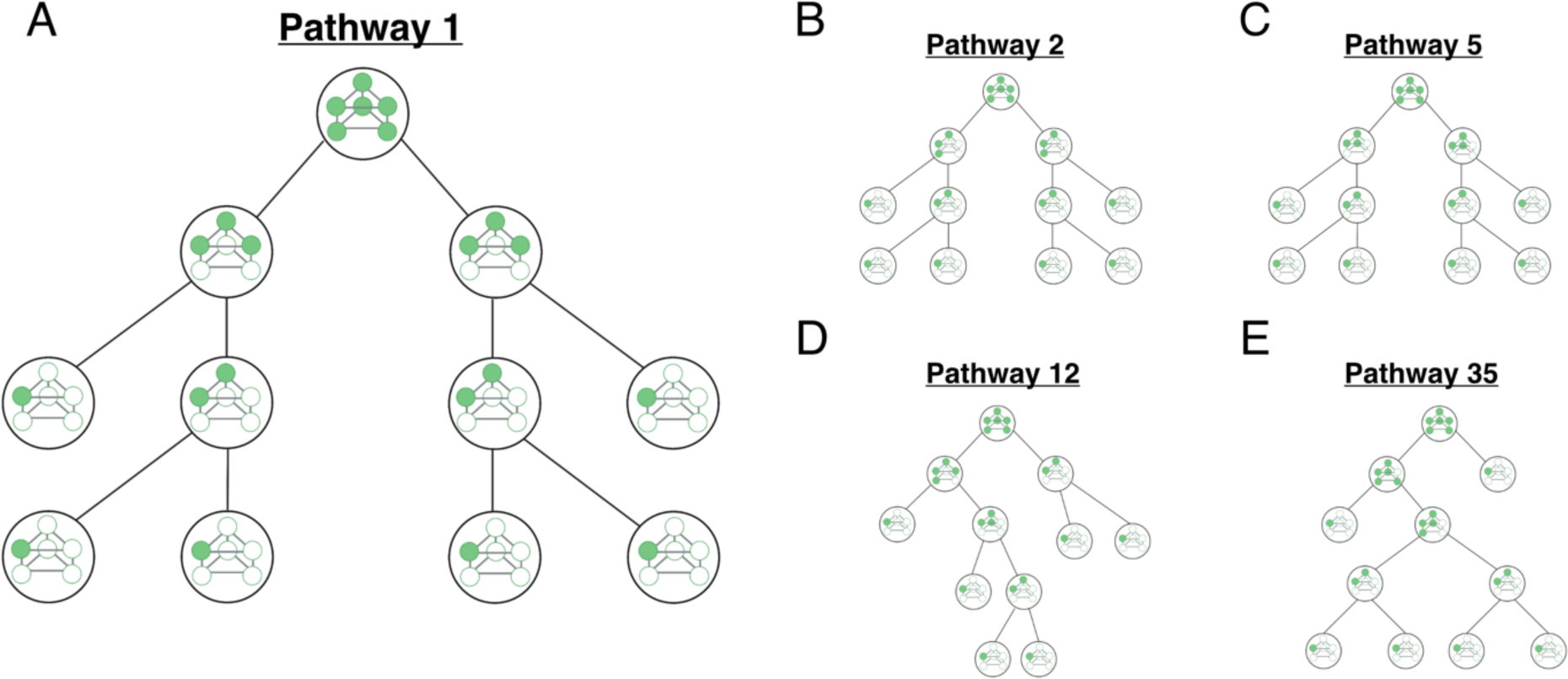
Example assembly pathways for a stacked trimer. Each panel shows an assembly pathway to form a stacked trimer. While we have enumerated all 46 pathways, 5 examples are shown here. The pathways begin at the bottom of the graphic and end with the stacked trimer at the top of the graphic. **(A)** Pathway 1 has a final step in which two trimeric rings bind to form a stacked trimer. **(B)** Pathway 2 is one pathway in which the final step is a reaction between two trimers (which in this case are not rings). **(C)** Pathway 5 represents an alternative to Pathway 2 in panel **B** with the same final step. **(D)** Pathway 12 is one pathway for which the final step is the reaction between a tetramer and a dimer to form the stacked trimer. **(E)** Pathway 35 is one pathway in with the final reaction is to add a single monomer to a pentamer to form the stacked trimer.

The pathway shown in **Figure 5A** represents one intuitive assembly pathway. Here, two monomers first combine to form a *within-ring* dimer; in other words, they bind to form one *K*_*d*,1_ interface. Another monomer then binds to form a complete ring. Two of these rings then dimerize with one another to form the fully-assembled structure (**Figure 5A**). While this is a very natural pathway, many other scenarios are possible. For instance, the pathways in **Figures 5B** and **5C** involve the formation of trimers that are not rings but can nonetheless combine into the fully-assembled stack trimer. **Figure 5D** shows a pathway where a dimer binds to a tetramer in the last step, and **Figure 5E** shows a case where monomers bind one-at-a-time.

It is straightforward to define a recursive algorithm that can find all the assembly pathways that are possible for a given structure. To do so, we start with the root node and consider all the possible “final reactions” that have the fully-assembled structure as their product. In the case of the stacked trimer, there are six such reactions, corresponding to six different ways that the final step in assembly can occur. Every one of these reactions has two reactants; these are the first two child nodes of the root node. For each child node, we can similarly enumerate all the unique reactions that have that species as their product; that is all the possible sets of child nodes for that particular node in the tree. We can continue this process recursively until we have only monomers as reactants, keeping track of each unique way of constructing each intermediate in the process. Using this approach, we enumerated all 46 possible assembly pathways for the stacked trimer; these are shown in **Supplemental Material Section 4, Figure S12**. The pathways in **Figure 5** are numbered according to the particular recursive scheme that we used to enumerate them. The code used for assembly pathway definitions is available at: https://github.com/llagunes-324B21/stackedTrimer_detModel.

### Pathway contributions reveal the origins of deadlock

We hypothesized that variation in the values of *K*_*d*,1_ and *K*_*d*,2_ could change how much any one of these 46 pathways “contributes” to the formation of the fully assembled stacked trimer. To do so, however, we must first mathematically define this notion of contribution. As described in **Supplemental Material Section 4.2**, we have enumerated all the possible binding and unbinding reactions that can occur in the assembly of the stacked trimer. Every binding reaction can be written 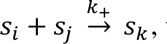 where *s*_*i*_ and *s*_j_ are the reactant species and *s*_*k*_ the product (these *s*’s are any species in **Figure 1C**). We call the set of all binding reactions *B* and the subset of those reactions that have species “*s*_*k*_” as their product *B*(*s*_*k*_). More formal definitions of a “reaction” and a “set of reactions” is given in **Supplemental Material Section 1.3**.

We define a pathway’s contribution to the formation of the fully stacked trimer in a recursive way as follows. First consider the fully assembled structure, Species “*s*_12_” in **Figure 1C**. All the different reactions that can produce this structure are collected in the subset *A*(*s*_12_). Take some reaction *r* ∈ *A*(*s*_12_), and define the *flux of the reaction r* as *F*(*r*). For instance, call the binding reaction *s*_1_ + *s*_11_ → *s*_12_ as “*r*_1_”: this is just the monomer binding to the pentamer to form the fully assembled stacked trimer, and clearly *r*_1_ ∈ *B*(*s*_12_). The flux of this reaction is just *F*(*r*_1_) = *k*_+_*x*_1_*x*_11_. Note that these fluxes are, in general, dependent on time, but as before we leave this dependence on time implicit. The expressions for the fluxes can also be more complicated, depending on things like stoichiometric coefficients; a full description of how to define any given flux is given in **Supplemental Material Section 1.5**. Finally, it is important to remember that a flux is not an *amount* or *concentration*, but rather *concentration per unit time*. For instance, we might have *F*(*r*_1_) = 1 *μ*M s^-1^, which means the reaction in question is producing 1 micromolar of the fully assembled structure every second.

Now that we have defined this flux, we can define the *total flux* of reactions that produce a given species *s*_*k*_ as *F*_*T*_(*s*_*k*_) ≡ ∑_*r*∈*B*(*s*__*k*)_ *F*(*r*). This just quantifies how much of that species is being formed per unit time, no matter which reaction is producing it. This finally allows us to define the critical quantity we need, which is the *relative flux of a reaction* 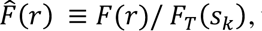 where *F*_*T*_(*s*_*k*_) is just the total flux of production of whatever species is the product of the reaction in question. This quantity can be naturally interpreted in the following way. Say we are considering the reaction *r*_1_ described above. Imagine we have a single new molecule of the fully assembled structure, which is the product of this reaction. 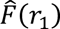 is just the probability that this molecule of *s*_12_ was produced by reaction *r*_1_.

By definition, every assembly pathway has some reaction *r* ∈ *B*(*s*_12_) as its “final step,” and there are only six such reactions (see the top of the panels in **Figure 6**). We can calculate the relative flux of this reaction, and that is the probability that all the pathways with that last step have produced any given “newly assembled” molecule of the fully assembled structure. To calculate the probability that an entire *pathway* is being used to form the fully-assembled structure, we now need to consider the two species that are combining to form the fully assembled structure. For instance, “Pathway 35” in **Figure 5E** has reaction *r*_1_ ∶ *s*_1_ + *s*_11_ → *s*_12_ as its last step. Since *s*_1_ is the monomer, we don’t need to consider how it is formed. Species *s*_11_, however, could be formed in several different ways, and Pathway 35 indicates that it is formed by the reaction *s*_1_ + *s*_8_ → *s*_11_, which we will call reaction “*r*_2_” for simplicity. Note that a fully assembled stacked trimer that uses Pathway 35 requires that both *r*_1_ AND *r*_2_ be used for the formation of the fully assembled structure; the probability that a stacked trimer is formed by these two reactions as the last and second-to-last reactions, respectively, is naturally 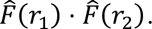

**Figure 6:**
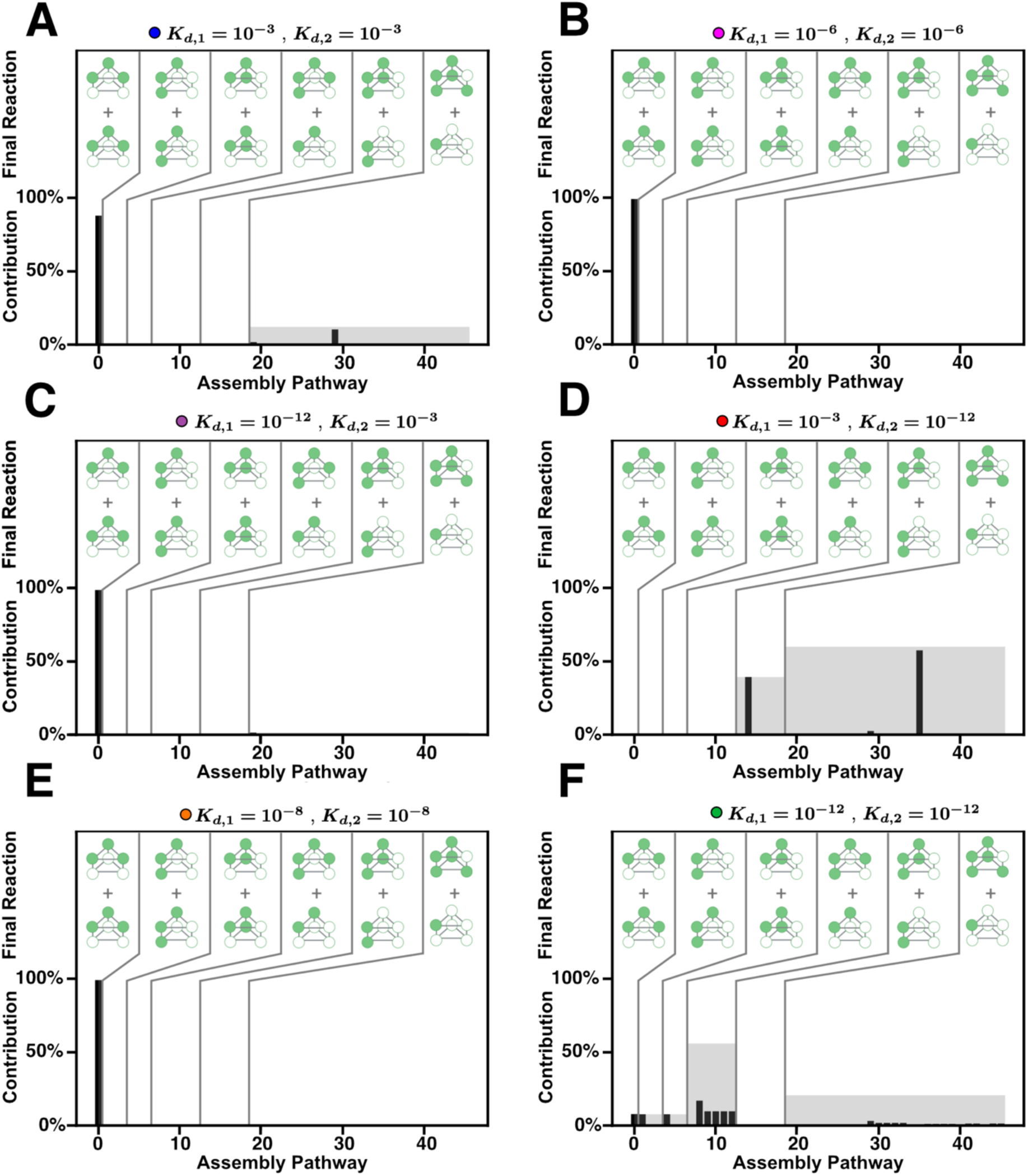
Assembly pathway contributions for different combinations of *K*_*d*,1_ and *K*_*d*,2_ *in vitro*. Each panel shows all the assembly pathways, enumerated as in Figure 4, and their calculated pathway contribution. Pathways are grouped based on the final assembly reaction as indicated by the graphics in the top of each graph labeled as “Final Reaction.” The black bar is the individual pathway contribution, and the gray region represents the sum of the pathway contributions for all the pathways in the group.

Any assembly pathway can be seen as a subset of binding reactions; in other words, a pathway *P*_*i*_ can be thought of as a subset of *B* (*P*_*i*_ ⊂ *B*). We can follow the recursive logic above and realize that the probability of a molecule of the stacked trimer forming *using* that pathway is just the product of the relative fluxes of all the reactions in the pathway. This leads us to define the *contribution of a pathway P*_*i*_ as: 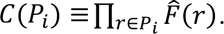 While the definition in this case may seem a bit cumbersome, the final outcome is quite straightforward. The larger the value of *C*(*P*_*i*_) for a given pathway, the higher the chance that a molecule of the fully assembled structure was formed according to the scenario described in the pathway. For instance, if *C*(*P*_1_) = 1, then the pathway in **Figure 5A** is the *only* way stacked trimer is being formed; alternatively, if *C*(*P*_1_) = 0, then the scenario in **Figure 5A** is not occurring at all. Note that this definition is similar to those used in previous work quantifying how different biochemical pathways contribute to enzyme flux (88). Further information on the definition and calculation of pathway contribution is given in the **Supplemental Material Section 4**.

In **Figure 6**, we calculate the pathway contribution for each pathway in the *in vitro* scenario after 24 hours with fixed binding affinity values. Here, pathways are grouped by their final association reaction, depicted on the top half of each contribution bar graph. There is only one pathway in which the final association step is the binding between two trimeric rings, Pathway 1. There are 3 pathways in which the final association step is the binding between two trimers that are not rings, 3 pathways with two trimers in a different orientation, 6 pathways with a *K*_*d*,1_-dimer binding to a tetramer, 6 pathways with a *K*_*d*,2_-dimer binding to a tetramer and finally there are 27 pathways where the final association step is the addition of a single monomer to a hexamer. In **Figure 6A**, we see that when both binding affinities *K*_*d*,1_ and *K*_*d*,2_ are weak, Pathway 1 contirbutes the most to the overall assembly of the stacked trimer, with only two pathways from the last group contributing less than 20%. This is likely due to the fact that the trimer is naturally much more stable than any of the dimers (15), so when the interactions are weak, the only fully assembled structures that can form rely on this relatively stable intermediate. We see a similar trend in **Figure 6B, C** and **E**, where one pathways contributes the most to the overall assembly of the stacked trimer. In **Figure 6D**, *K*_*d*,1_ is relatively weak and *K*_*d*,2_ is very strong; here we see that two pathways contribute almost equally, which are the pathways in which the final association reaction is between a *K*_*d*,2_-dimer binding to a tetramer or a single monomer addition to the hexamer. Since *K*_*d*,2_ is very strong in this scenario, this suggests that many of the monomers may be “stuck” in that a *K*_*d*,2_-dimer and not able to bind with other intermediates formed, which leads to deadlock. More interestingly, when both binding affinities *K*_*d*,1_ and *K*_*d*,2_ are very strong, **Figure 6F** shows many pathways contributing to the overall yield of the stacked trimer. This shows that when both binding affinities are strong, many of the possible pathways are taking place and consuming monomers. Since these pathways rely on different intermediate structures that may not be compatible, this leads to high levels of deadlock. In other words, having very strong afinities for both interactions in the structure leads to broad utlization of many pathways and deadlock, whereas having one of the interfaces significantly weaker induces a more hierarchical scenario where only one or a few pathways contribute (**Figure 6**).

We see similar results for the *in vivo* model. Note that in **Figure 6** we are considering the relative fluxes at a particular time; in **Figure 7** we are considering pathway contributions at steady state for the *in vivo* model. When both binding affinities *K*_*d*,1_ and *K*_*d*,2_ are weak, **Figure 7A** shows that few pathways contribute the most to the assembly of the stacked trimer. Interestingly, **Figure 7B-D** show that more pathways contribute to the stacked trimer assembly than in the *in vitro* case when one of the interactions is significantly weaker than the other. This is likely due to the constant synthesis of monomers allowing more pathways to contribute. Regardless, in all of these cases only a small number of pathways contributge siginficantly to the formation of the stacked trimer. However, when both binding affinities *K*_*d*,1_ and *K*_*d*,2_ are strong, **Figures 7E** and **F** show that many pathways contribute. As in the *in vitro* case, this leads to higher steady-state levels of incompatible intermediates and thus a lower overall flux of stacked trimer production. This analysis of the contribution of assembly pathways indicates that specific patterns of affinity have evolved to enforce a more “hierarcichal” assembly process that relys on only a few pathways (15).

**Figure 7:**
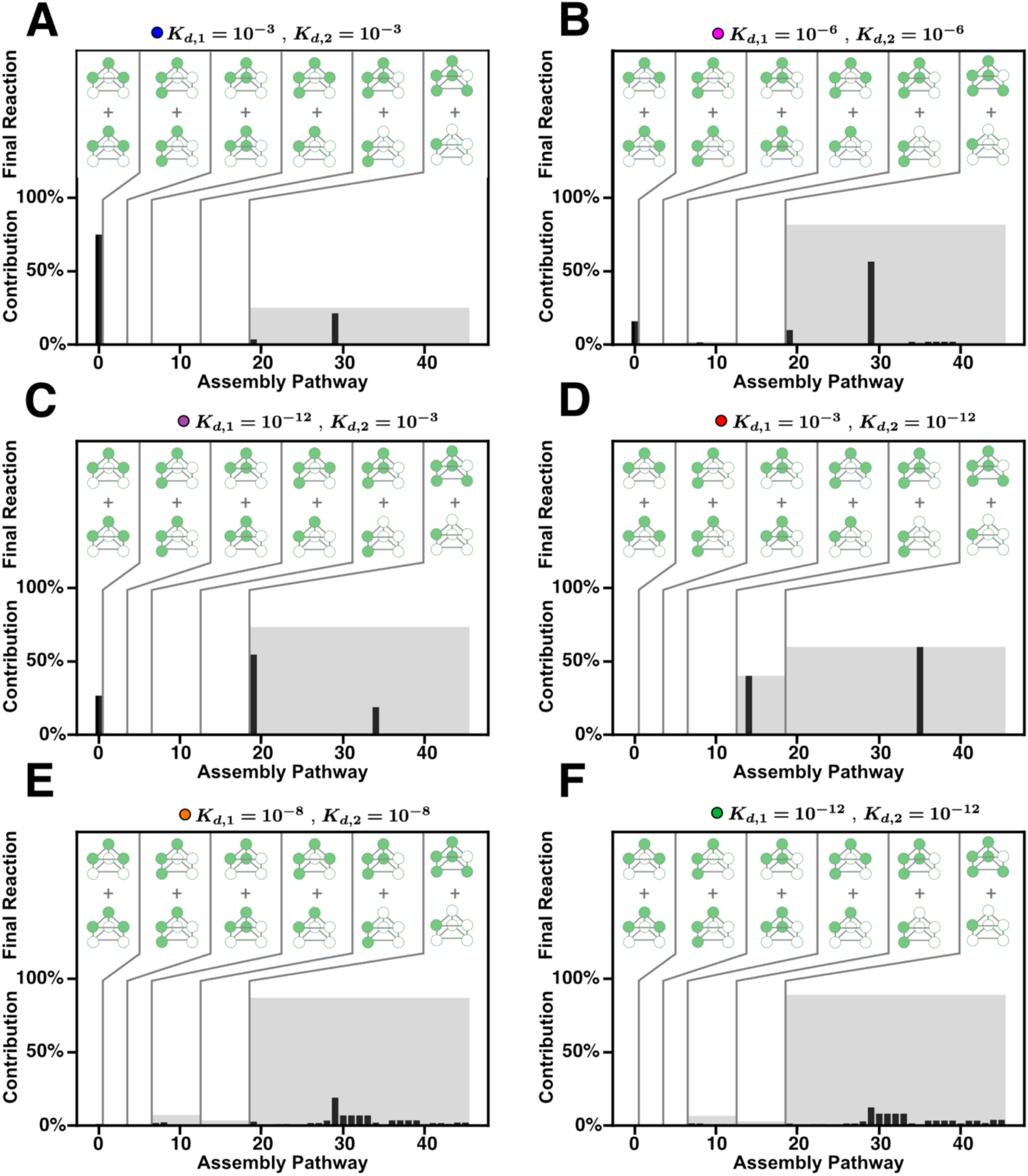
Assembly pathway contributions for different combinations of *K*_*d*,1_ and *K*_*d*,2_ *in vivo*. Each panel shows all the assembly pathways, enumerated as in Figure 4, and their calculated pathway contribution. Pathways are grouped based on the final assembly reaction as indicated by the graphics in the top of each graph, labeled as “Final Reaction.” The black bar is the individual pathway contribution, and the gray region represents the sum of the pathway contributions for all the pathways in the group.

## Discussion

Almost every cellular process relies in some way or another on the function of a variety of different macromolecular machines. Understanding how these machines are assembled is thus critical to our overall understanding of their function and ultimately the regulation of a wide variety of cellular activities. In recent years, new work has emerged that focuses on the design of synthetic protein complexes for applications such as drug delivery, gene therapy and biofuel production (12–14). Understanding the assembly of natural complexes can also provide potential “design principles” for the design of new synthetic machines in order to optimize self-assembly. This will also aid in advancing biotechnological applications and facilitate the integration of natural and synthetic systems for innovative solutions in medicine, biotechnology, and sustainable energy production.

Many experimental studies have described the assembly processes of various molecular machines (31–37, 89). Although these experiments have expanded our understanding of the assembly of specific structures, including that of stacked ring-like structures, many details of the assembly process remain elusive. For instance, this experimental work does not address how assembly is optimized to avoid phenomena like kinetic trapping. Mathematical models offer an alternative approach to studying assembly, allowing us to describe the assembly process in great detail and understand how mechanisms of assembly have evolved to improve assembly efficiency (15). Specifically, both on-pathway and off-pathway kinetic trapping have been observed in a wide variety of mathematical and biophysical models of assembly processes, and previous work has suggested that structures, particularly ring-like structures, have likely evolved to avoid on-pathway kinetic trapping (in other words, deadlock).

In this work, we expanded on previous mathematical frameworks to understand the mechanisms of stacked ring assembly. We found that the phenomenon of deadlock can be much more severe in the case of stacked rings than had been observed for simple ring-like structures. In particular, deadlock can result in much lower assembly yields and take much longer to resolve for stacked rings than it does for single rings. As in the case of single rings, the severity of deadlock depends on the strength of the interactions between the subunits in the ring; if all the interactions are strong, deadlock can severely reduce assembly yields on biologically realistic timescales. This suggested an evolutionary pressure to avoid cases where all the interfaces in a stacked ring have high affinity. Extensive analysis of solved structures of stacked rings suggests that evolution has indeed avoided this particular scenario. Interestingly, it is common for the design pipelines of synthetic macromolecular assemblies to minimize the binding energies of *every* interface. Our work suggests that this may be a highly suboptimal approach to the design of structures that self-assemble efficiently, a factor that may in part explain the high failure rate of these designs. Our work thus reveals simple principles that may be of use to future efforts in protein design.

We also uses our models to explore the notion of an assembly pathway in greater detail. Many experimental studies discuss the notion of “assembly pathways” (90–92) but have no *precise* definition of what an assembly pathway actually is. Here, we provide a more formal, mathematical definition of an assembly pathway, which we expect will provide a structured framework for understanding the intricate process of macromolecular assembly and enable precise analysis, prediction and manipulation of these critical biological mechanisms. Application of this framework to our models revealed exactly *why* having strong interactions in these structures generates deadlock. There are many different scenarios whereby a set of monomers can proceed towards the fully-assembled structure. When interactions are strong, the system simultaneously proceeds down all these paths, depleting resources and leading to a situation where a large number of incompatible intermediates are formed (i.e. deadlock). When some of the interactions are weak, however, only a small subset of these pathways are employed, leading to a naturally hierarchical assembly process that is much more resistant to kinetic trapping.

Future research will be necessary to further link the results of models like these to experimental studies of macromolecular machines. For instance, the proteasome is a barrel-like structure made up of four heptameric rings in an α_7_β_7_β_7_α_7_ stoichiometry that is conserved across all kingdoms of life. In archaea, there is experimental evidence that the outer α_7_ rings assemble first and then the β subunits bind to the α_7_ ring (31, 93). However, in bacteria the outer α_7_ rings cannot assemble on their own, and instead, both α and β are necessary for self-assembly *in vitro* (2, 31). This has led to the proposal that these two different proteasomes utilize different assembly pathways, but detailed predictions on how these scenarios might lead to experimentally-testable hypotheses is lacking. Thus, in addition to a need for more detailed experimental work, there is also a critical need for the development of mathematical and biophysical models of these more complex structures. These models will prove critical to our understanding of the design principles of macromolecular machines and the application of those principles to synthetic biology.

## Notes

### Competing Interest Statement

The authors have declared no competing interest.

## References

1. Yuan, S., G. Zhou, and G. Xu. 2023. Translation machinery: the basis of translational control. Journal of Genetics and Genomics.

2. Budenholzer, L., C.L. Cheng, Y. Li, and M. Hochstrasser. 2017. Proteasome Structure and Assembly. J Mol Biol. 429:3500–3524.

3. Stan, G., G.H. Lorimer, and D. Thirumalai. 2022. Friends in need: How chaperonins recognize and remodel proteins that require folding assistance. Front Mol Biosci. 9.

4. Prywes, N., N.R. Phillips, O.T. Tuck, L.E. Valentin-Alvarado, and D.F. Savage. 2023. Annual Review of Biochemistry Rubisco Function, Evolution, and Engineering..

5. Zhou, A., A. Rohou, D.G. Schep, J. V Bason, M.G. Montgomery, J.E. Walker, N. Grigorieff, and J.L. Rubinstein. 2015. Structure and conformational states of the bovine mitochondrial ATP synthase by cryo-EM. Elife.

6. Nogales, E. 2000. Structural Insights into Microtubule Function. Annual Reviews in Biochemistry. 69:277–302.

7. Chen, Y., Y. Zhang, and X. Guo. 2017. Proteasome dysregulation in human cancer: implications for clinical therapies. Cancer and Metastasis Reviews. 36:703–716.

8. Fhu, C.W., and A. Ali. 2021. Dysregulation of the ubiquitin proteasome system in human malignancies: A window for therapeutic intervention. Cancers (Basel). 13.

9. Zheng, Q., T. Huang, L. Zhang, Y. Zhou, H. Luo, H. Xu, and X. Wang. 2016. Dysregulation of ubiquitin-proteasome system in neurodegenerative diseases. Front Aging Neurosci. 8.

10. Versari, D., J. Herrmann, M. Gössl, D. Mannheim, K. Sattler, F.B. Meyer, L.O. Lerman, and A. Lerman. 2006. Dysregulation of the ubiquitin-proteasome system in human carotid atherosclerosis. Arterioscler Thromb Vasc Biol. 26:2132–2139.

11. Cao, J., M.B. Zhong, C.A. Toro, L. Zhang, and D. Cai. 2019. Endo-lysosomal pathway and ubiquitin-proteasome system dysfunction in Alzheimer’s disease pathogenesis. Neurosci Lett. 703:68–78.

12. Smith, L.C., J. Duguid, M.S. Wadhwa, M.J. Logan, C. Hsuan Tung, V. Edwards, and J.T. Sparrow. 1998. Synthetic peptide-based DNA complexes for nonviral gene delivery..

13. Chao, J., H. Liu, S. Su, L. Wang, W. Huang, and C. Fan. 2014. Structural DNA nanotechnology for intelligent drug delivery. Small. 10:4626–4635.

14. Wagner, H.J., W. Weber, and M. Fussenegger. 2021. Synthetic Biology: Emerging Concepts to Design and Advance Adeno-Associated Viral Vectors for Gene Therapy. Advanced Science. 8.

15. Deeds, E.J., J.A. Bachman, W. Fontana, W.F. Designed, and J.A.B. Performed. 2012. Optimizing ring assembly reveals the strength of weak interactions. PNAS. 109.

16. Levy, E.D., J.B. Pereira-Leal, C. Chothia, and S.A. Teichmann. 2006. 3D complex: A structural classification of protein complexes. PLoS Comput Biol. 2:1395–1406.

17. Levy, E.D., E.B. Erba, C. V. Robinson, and S.A. Teichmann. 2008. Assembly reflects evolution of protein complexes. Nature. 453:1262–1265.

18. Almassy, R.J., C.A. Janson, J.R. Hamlin, N.-H. Xuong, and D. Eisenberg. 1986. Novel subunit-subunit interactions in the structure of glutamine synthase. Nature. 323:304–309.

19. Eisenberg, D., H.S. Gill, G.M.U. P£uegl, and S.H. Rotstein. 2000. Structure-function relationships of glutamine synthetases 1. 122–145.

20. Schormann, N., K.L. Hayden, P. Lee, S. Banerjee, and D. Chattopadhyay. 2019. An overview of structure, function, and regulation of pyruvate kinases. Protein Science. 28:1771–1784.

21. Pattanayek, R., J. Wang, T. Mori, Y. Xu, C.H. Johnson, and M. Egli. 2004. Visualizing a Circadian Clock Protein: Crystal Structure of KaiC and Functional Insights. Mol Cell. 15:375–388.

22. Vale, R.D. 2000. Mini-Review AAA Proteins: Lords of the Ring..

23. Lupas, A.N., and J. Martin. 2002. AAA proteins. Current Opinion in Structural Biology. 12:746–753.

24. Marques, A.J., R. Palanimurugan, A.C. Mafias, P.C. Ramos, and R.J. Dohmen. 2009. Catalytic mechanism and assembly of the proteasome. Chem Rev. 109:1509–1536.

25. van Wijk, S.J.L., and H.T.M. Timmers. 2010. The family of ubiquitin-conjugating enzymes (E2s): deciding between life and death of proteins. The FASEB Journal. 24:981–993.

26. Dokland, T. 2000. Freedom and restraint: themes in virus capsid assembly. Structure. 8:157–162.

27. Hagan, M.F., and D. Chandler. 2006. Dynamic pathways for viral capsid assembly. Biophys J. 91:42–54.

28. Selivanovitch, E., and T. Douglas. 2019. Virus capsid assembly across different length scales inspire the development of virus-based biomaterials. Curr Opin Virol. 36:38–46.

29. Zhou, W., T. Šmidlehner, and R. Jerala. 2020. Synthetic biology principles for the design of protein with novel structures and functions. FEBS Lett. 594:2199–2212.

30. Hamley, I.W. 2019. Protein Assemblies: Nature-Inspired and Designed Nanostructures. Biomacromolecules. 20:1829–1848.

31. Zühl, F., E. Seemuller, R. Golbik, and W. Baumeister. 1997. Dissecting the assembly pathway of the 20S proteasome. FEBS Lett. 418:189–194.

32. Witt, S., Y. Do Kwon, M. Sharon, K. Felderer, M. Beuttler, C. V. Robinson, W. Baumeister, and B.K. Jap. 2006. Proteasome Assembly Triggers a Switch Required for Active-Site Maturation. Structure. 14:1179–1188.

33. Bernstein, K.A., J.E.G. Gallagher, B.M. Mitchell, S. Granneman, and S.J. Baserga. 2004. The small-subunit processome is a ribosome assembly intermediate. Eukaryot Cell. 3:1619–1626.

34. Bogenhagen, D.F., A.G. Ostermeyer-Fay, J.D. Haley, and M. Garcia-Diaz. 2018. Kinetics and Mechanism of Mammalian Mitochondrial Ribosome Assembly. Cell Rep. 22:1935–1944.

35. Voziyan, P.A., and M.T. Fisher. 2002. Polyols induce ATP-independent folding of GroEL-bound bacterial glutamine synthetase. Arch Biochem Biophys. 397:293–297.

36. O’Neil, P.T., A.J. Machen, B.C. Deatherage, C. Trecazzi, A. Tischer, V.R. Machha, M.T. Auton, M.R. Baldwin, T.A. White, and M.T. Fisher. 2018. The Chaperonin GroEL: A Versatile Tool for Applied Biotechnology Platforms. Front Mol Biosci. 5.

37. Ryabova, N.A., V. V. Marchenkov, S.Y. Marchenkova, N. V. Kotova, and G. V. Semisotnov. 2013. Molecular chaperone GroEL/ES: Unfolding and refolding processes. Biochemistry (Moscow). 78:1405–1414.

38. Kumar, M.S., and R. Schwartz. 2010. A parameter estimation technique for stochastic self-assembly systems and its application to human papillomavirus self-assembly. Phys Biol. 7.

39. Thomas, M., and R. Schwartz. 2018. A method for efficient Bayesian optimization of self-assembly systems from scattering data. BMC Syst Biol. 12.

40. Thomas, M., and R. Schwartz. 2017. Quantitative computational models of molecular self-assembly in systems biology. Phys Biol. 14.

41. Nguyen, H.D., V.S. Reddy, and C.L. Brooks. 2007. Deciphering the kinetic mechanism of spontaneous self-assembly of icosahedral capsids. Nano Lett. 7:338–344.

42. Mannige, R. V, and C.L.I. Brooks. 2009. Geometric considerations in virus capsid size specificity, auxiliary requirements, and buckling. 8531–8536.

43. Mannige, R. V., and C.L. Brooks. 2010. Periodic table of virus capsids: Implications for natural selection and design. PLoS One. 5.

44. Grime, J.M.A., J.F. Dama, B.K. Ganser-Pornillos, C.L. Woodward, G.J. Jensen, M. Yeager, and G.A. Voth. 2016. Coarse-grained simulation reveals key features of HIV-1 capsid self-assembly. Nat Commun. 7.

45. Banerjee, P., and G.A. Voth. 2023. Conformational transitions of the HIV-1 Gag polyprotein upon multimerization and gRNA binding. Biophys J.

46. Williamson, J.R. 2008. Biophysical studies of bacterial ribosome assembly. Curr Opin Struct Biol. 18:299–304.

47. Burton, B., M.T. Zimmermann, R.L. Jernigan, and Y. Wang. 2012. A computational investigation on the connection between dynamics properties of ribosomal proteins and ribosome assembly. PLoS Comput Biol. 8.

48. Perilla, J.R., B.C. Goh, C.K. Cassidy, B. Liu, R.C. Bernardi, T. Rudack, H. Yu, Z. Wu, and K. Schulten. 2015. Molecular dynamics simulations of large macromolecular complexes. Curr Opin Struct Biol. 31:64–74.

49. Goyal, A., K. Muthu, M. Panneerselvam, A.K. Pole, and K. Ramadas. 2011. Molecular dynamics simulation of the Staphylococcus aureus YsxC protein: Molecular insights into ribosome assembly and allosteric inhibition of the protein. J Mol Model. 17:3129–3149.

50. Suppahia, A., P. Itagi, A. Burris, F.M.G. Kim, A. Vontz, A. Kante, S. Kim, W. Im, E.J. Deeds, and J. Roelofs. 2020. Cooperativity in Proteasome Core Particle Maturation. iScience. 23.

51. Sharon, M., S. Witt, E. Glasmacher, W. Baumeister, and C. V. Robinson. 2007. Mass spectrometry reveals the missing links in the assembly pathway of the bacterial 20 S proteasome. Journal of Biological Chemistry. 282:18448–18457.

52. Vakser, I.A., and E.J. Deeds. 2019. Computational approaches to macromolecular interactions in the cell. Curr Opin Struct Biol. 55:59–65.

53. Vakser, I.A., S. Grudinin, N.W. Jenkins, P.J. Kundrotas, and E.J. Deeds. 2022. Docking-based long timescale simulation of cell-size protein systems at atomic resolution..

54. Tinberg, C.E., S.D. Khare, J. Dou, L. Doyle, J.W. Nelson, A. Schena, W. Jankowski, C.G. Kalodimos, K. Johnsson, B.L. Stoddard, and D. Baker. 2013. Computational design of ligand-binding proteins with high affinity and selectivity. Nature. 501:212–216.

55. Allison, B., S. Combs, S. DeLuca, G. Lemmon, L. Mizoue, and J. Meiler. 2014. Computational design of protein-small molecule interfaces. J Struct Biol. 185:193–202.

56. Kosugi, T., T. Iida, M. Tanabe, R. Iino, and N. Koga. 2023. Design of allosteric sites into rotary motor V1-ATPase by restoring lost function of pseudo-active sites. Nat Chem. 15:1591–1598.

57. Miller, J.E., Y. Srinivasan, N.P. Dharmaraj, A. Liu, P.L. Nguyen, S.D. Taylor, and T.O. Yeates. 2022. Designing Protease-Triggered Protein Cages. J Am Chem Soc. 144:12681–12689.

58. Bale, J.B., R.U. Park, Y. Liu, S. Gonen, T. Gonen, D. Cascio, N.P. King, T.O. Yeates, and D. Baker. 2015. Structure of a designed tetrahedral protein assembly variant engineered to have improved soluble expression. Protein Science. 24:1695–1701.

59. Pfoh, R., E.F. Pai, and V. Saridakis. 2015. Nicotinamide mononucleotide adenylyltransferase displays alternate binding modes for nicotinamide nucleotides. Acta Crystallogr D Biol Crystallogr. 71:2032–2039.

60. Ho, W.W., H. Li, S. Eakanunkul, Y. Tong, A. Wilks, M. Guo, and T.L. Poulos. 2007. Holo- and apo-bound structures of bacterial periplasmic heme-binding proteins. Journal of Biological Chemistry. 282:35796–35802.

61. Hadler, K.S., E.A. Tanifum, S.H.C. Yip, N. Mitić, L.W. Guddat, C.J. Jackson, L.R. Gahan, K. Nguyen, P.D. Carr, D.L. Ollis, A.C. Hengge, J.A. Larrabee, and G. Schenk. 2008. Substrate-promoted formation of a catalytically competent binuclear center and regulation of reactivity in a glycerophosphodiesterase from Enterobacter aerogenes. J Am Chem Soc. 130:14129–14138.

62. Zhang, T., and R. Schwartz. 2006. Simulation study of the contribution of oligomer/oligomer binding to capsid assembly kinetics. Biophys J. 90:57–64.

63. Xie, L., G.R. Smith, X. Feng, and R. Schwartz. 2012. Surveying capsid assembly pathways through simulation-based data fitting. Biophys J. 103:1545–1554.

64. Saiz, L., and J.M.G. Vilar. 2006. Stochastic dynamics of macromolecular-assembly networks. Mol Syst Biol. 2.

65. Duke, T.A.J., N. Le Novère, and D. Bray. 2001. Conformational spread in a ring of proteins: A stochastic approach to allostery. J Mol Biol. 308:541–553.

66. Camacho, C.J., S.R. Kimura, C. Delisi, and S. Vajda. 2000. Kinetics of Desolvation-Mediated Protein-Protein Binding..

67. Tait, S.W.G., and D.R. Green. 2010. Mitochondria and cell death: Outer membrane permeabilization and beyond. Nat Rev Mol Cell Biol. 11:621–632.

68. Shiozaki, E.N., J. Chai, and Y. Shi. 2001. Oligomerization and activation of caspase-9, induced by Apaf-1 CARD..

69. Teng, X., and J.M. Hardwick. 2010. The apoptosome at high resolution. Cell. 141:402–404.

70. Qi, H., Y. Jiang, Z. Yin, K. Jiang, L. Li, and J. Shuai. 2018. Optimal pathways for the assembly of the Apaf-1·cytochrome: C complex into apoptosome. Physical Chemistry Chemical Physics. 20:1964–1973.

71. Mangan, S., and U. Alon. 2003. Structure and function of the feed-forward loop network motif..

72. Shtilerman, M., G.H. Lorimer, and S.W. Englander. 2012. Chaperonin Function: Folding by Forced Unfolding. Science (1979). 284:822–825.

73. Klinge, S., and J.L. Woolford. 2019. Ribosome assembly coming into focus. Nat Rev Mol Cell Biol. 20:116–131.

74. Kelman, Z., and M. O’donnell. 1995. DNA POLYMERASE III HOLOENZYME: Structure and Function of a Chromosomal Replicating Machine..

75. Farnung, L., and S.M. Vos. 2022. Assembly of RNA polymerase II transcription initiation complexes. Curr Opin Struct Biol. 73.

76. Maurizi, M.R. 1998. Proteasome assembly: Bitting the hand… Current Biology. 8:453–456.

77. Jonckheere, A.I., J.A.M. Smeitink, and R.J.T. Rodenburg. 2012. Mitochondrial ATP synthase: Architecture, function and pathology. J Inherit Metab Dis. 35:211–225.

78. Pagliano, C., G. Saracco, and J. Barber. 2013. Structural, functional and auxiliary proteins of photosystem II. Photosynth Res. 116:167–188.

79. Lynch, M., and G.K. Marinov. 2015. The bioenergetic costs of a gene. Proc Natl Acad Sci U S A. 112:15690–15695.

80. Schavemaker, P.E., and M. Lynch. 2022. Flagellar energy costs across the tree of life. Elife. 11.

81. Horton, N., and M. Lewis. 1992. Calculation of the free energy of association for protein complexes. Protein Science. 1:169–181.

82. Bougouffa, S., and J. Warwicker. 2008. Volume-based solvation models out-perform area-based models in combined studies of wild-type and mutated protein-protein interfaces. BMC Bioinformatics. 9.

83. Kelly, J.W. 1998. The alternative conformations of amyloidogenic proteins and their multi-step assembly pathways. Curr Opin Struct Biol. 8:101–106.

84. Neuenkirchen, N., A. Chari, and U. Fischer. 2008. Deciphering the assembly pathway of Sm-class U snRNPs. FEBS Lett. 582:1997–2003.

85. Yoda, M., T. Kawamata, Z. Paroo, X. Ye, S. Iwasaki, Q. Liu, and Y. Tomari. 2010. ATP-dependent human RISC assembly pathways. Nat Struct Mol Biol. 17:17–24.

86. Mulder, A.M., C. Yoshioka, A.H. Beck, A.E. Bunner, R.A. Milligan, C.S. Potter, B. Carragher, and J.R. Williamson. 2010. Visualizing ribosome biogenesis: Parallel assembly pathways for the 30S subunit. Science (1979). 330:673–677.

87. Jacobs, W.M., A. Reinhardt, and D. Frenkel. 2015. Rational design of self-assembly pathways for complex multicomponent structures. Proc Natl Acad Sci U S A. 112:6313–6318.

88. Shockley, E.M., C.A. Rouzer, L.J. Marnett, E.J. Deeds, and C.F. Lopez. 2019. Signal integration and information transfer in an allosterically regulated network. NPJ Syst Biol Appl. 5.

89. Chari, A., and U. Fischer. 2010. Cellular strategies for the assembly of molecular machines. Trends Biochem Sci. 35:676–683.

90. Ligas, J., E. Pineau, R. Bock, M.A. Huynen, and E.H. Meyer. 2019. The assembly pathway of complex I in Arabidopsis thaliana. Plant Journal. 97:447–459.

91. Gallastegui, N., and M. Groll. 2010. The 26S proteasome: assembly and function of a destructive machine. Trends Biochem Sci. 35:634–642.

92. Goehring, N.W., and J. Beckwith. 2005. Diverse paths to midcell: Assembly of the bacterial cell division machinery. Current Biology. 15.

93. Zwickl, P., J. Kleinz, and W. Baumeister. 1994. Critical elements in proteasome assembly. Structural Biology. 1.

94. Briggs, K. 2018. Disrupted Pathways: Generating Tunable Macromolecular Assembly Pathways. The University of Kansas.

