## Supplemental Appendix for "Modeling reveals the strength of weak interactions in stacked ring assembly"

Manuscript submitted to **Biophysical** *Journal*
Article

### Modeling reveals the strength of weak interactions in stacked ring assembly
### Supporting Information Appendix

Leonila Lagunes[1,2], Koan Briggs[3], Paige Martin-Holder[4], Zaikun Xu[5], Dustin Maurer[5], Karim Ghabra[6], and Eric J. Deeds[1,2,5]

[1]Department of Integrative Biology and Physiology, UCLA
[2]Institute for Quantitative and Computational Biosciences, UCLA
[3]Department of Physics, University of Kansas
[4]Department of Molecular Immunology, Microbiology and Genetics, UCLA
[5]Center for Computational Biology, University of Kansas
[6]Computational and Systems Biology IDP, UCLA
[a]Main Street Data, Kansas City, MI, USA
[b]Los Angeles, CA, USA
[c]Astrivis, Zurich, Switzerland
[d]Consultant, Vancouver, Washington

#### 1 CONSTRUCTING A MODEL OF STACKED RING ASSEMBLY

##### 1.1 Interfaces in the stacked trimer

To construct a model of stacked ring assembly, we need to have an understanding of the interfaces within the monomers that bind to form the stacked ring with two 3-membered homomeric rings. As discussed in the main text, the monomeric subunits (species $S_1$) are identical and have three distinct interfaces: the left and right "*within-ring*" interfaces, and the "*between ring*" interface. The left and right interfaces each create a bond between two monomers to form an intra-ring dimer (species $S_3$), which we call the "*within-ring interaction*", shown in **Figure S1**. The other interface forms the bonds between the top and bottom monomers forming an inter-ring dimer (species $S_2$) which we call the "*between-ring interaction*," shown in **Figure S1**. The $K_d$ values for these two interfaces are defined in the standard way based on the association and dissociation rates of the

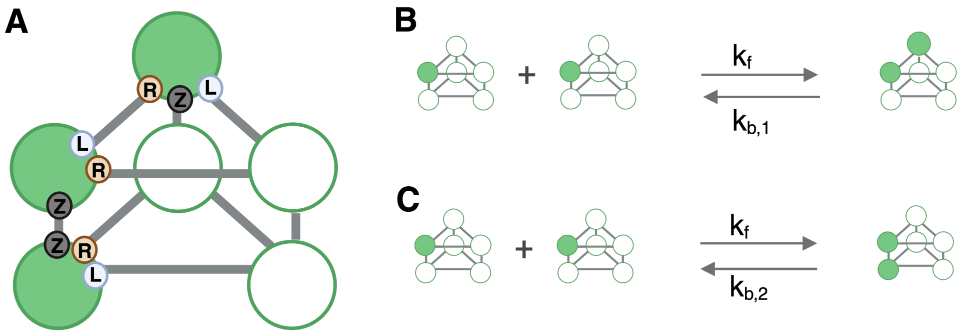

**Figure S1: Binding affinity reactions. (A)** Interfaces on $S_1$ monomer and it binding to a different monomer on a stacked ring. The **L** represents the "Left" interface of the monomer, the **R** represents the "Right" and the **Z** represents the opposite interface for binding between rings. Associated chemical reactions for (**B**) $K_{D,1}$ and (**C**) $K_{D,2}$. The monomers $S_1$ are depicted as the reactants and the two types of dimers $S_2$ and $S_3$ as the product. Following the same format as the main text, the filled in circles represent a present monomer and the unfilled circles represent an available location for a new monomer. Figure made with BioRender.

chemical reactions that that involve the formation of a single non-covalent interaction in a reversible manner; both chemical reactions are listed below:

$$S_1 + S_1 \underset{k_{b,1}}{\overset{k_f}{\rightleftharpoons}} S_3 \tag{1}$$

$$S_1 + S_1 \underset{k_{b,2}}{\overset{k_f}{\rightleftharpoons}} S_2 \tag{2}$$

For each chemical reaction above, we assume the forward reaction rate $k_\mathrm{f}$ to be equal and the backward as $k_{\mathrm{b},1}$ and $k_{\mathrm{b},2}$, respectively. Similar to the work done in [3], the standard free energy of formation of these interaction is related in the usual way to the dissociation constant $k_{\mathrm{b},i}$. Thus, the binding affinities for each reaction is:

$$K_{D,1} = \frac{k_\mathrm{f}}{k_{\mathrm{b},1}} \tag{3}$$

$$K_{D,2} = \frac{k_\mathrm{f}}{k_{\mathrm{b},2}} \tag{4}$$

In our model of stacked trimer assembly we take into account essential intermediates that are generated prior to the stacked timer. This determines how we write our ordinary differential equations. In this work, we used chemical reaction network theory to develop our ODEs similar to the work done in [3] where we account explicitly for combinatorial factors of a reaction rather than absorbing them into an single rate constant.

#### 1.2 Enumerating chemical species

To determine all of the intermediates that are assembled during stacked trimer assembly, we enumerated all the possible species that can be assembled from a monomer $S_1$ in our model (see **Figure 1C**). In enumerating these species, it is helpful to view any intermediate as a sub-structure or sub-graph of the fully assembled structure. In this approach, we have two kinds of nodes: the "filled in" nodes (dark green in **Figures S**1 and 2) represent subunits that are present in the structure, while the empty circles represent structures that are absent. Using this convention, we can represent the monomer (i.e. a single subunit) in six different ways **Figure S1A** corresponding to the rotational and mirror symmetry of the molecule. Since all of these are the same *chemical* species, we represent this as a single species $S_1$ (**Figure 1C**).

To enumerate the remainder of the species, we start with the monomer and add additional monomers sequentially. For instance, starting with the monomer $S_1$, we can add another monomer in one of three positions; to the left of that monomer in the top ring, to the right of the monomer in the top ring, and directly underneath the monomer. This results in two distinct types of dimers; in the first type of dimer, we form a single $K_{D,1}$ interaction (**Figure S2B**). While there are again six different ways to

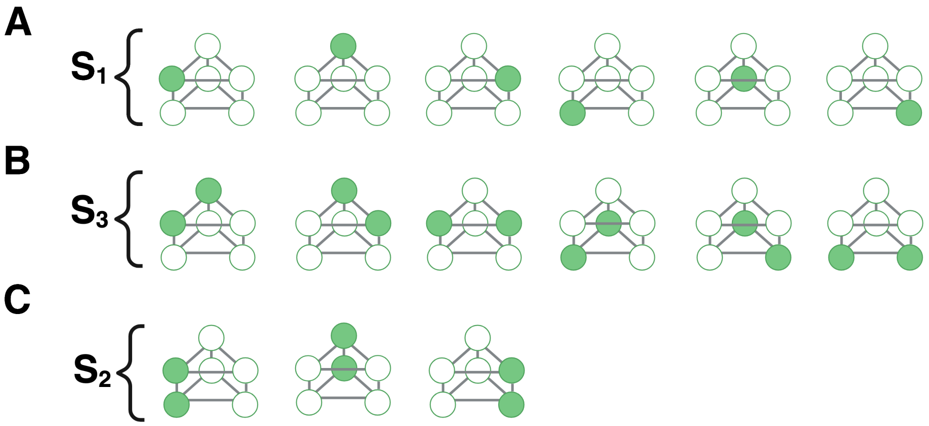

**Figure S2: Species consolidations.** Each structure can be consolidated into a single species based on symmetry. **(A)** The monomer can exist at any of the six available locations on the stacked trimer and they are all considered to be species $S_1$. **(B)** The $K_{D,1}$ dimer species $S_3$ is can be formed through any of the arrangements listed. Similarly with **(C)** the $K_{D,2}$ dimer species $S_2$. Figure made with BioRender.

represent this "$K_{D,1}$ dimer" (species $S_2$), we choose a single representative visualization for convenience (**Figure 1C**). If we add the monomer in the position directly across the ring from the other monomer, we form the "$K_{D,2}$ dimer" species $S_3$. Note that this structure also has six possible representations, but only 3 are shown in **Figure S2C** because the mirror image variants are visually equivalent to those shown in the figure.

We can enumerate all the trimers in much the same way. We start with the two dimer species $S_2$ and $S_3$; any trimeric species can be obtained by adding a monomer to each of these dimers. So we add a monomer in every "available" slot on each of the two dimers, which results in a set of species. Some of these are equivalent to one another because of the available symmetry operations (**Figure S2**); we collect those together into the set of unique trimer species. To generate the tetramers, we start with the resulting trimers and again add monomers in every available slot. Iterating this process results in the 12 unique chemical species shown in **Figure 1C** in the main text.

#### 1.3 Enumerating chemical reactions

To assemble our chemical reaction network of stacked trimer assembly, we enumerated all the possible reactions that can occur between species in this model (see **Figure 1C** in the main text). Formally speaking, a chemical reaction represents either a way of combining two species to make a third (this is an *association reaction*) or a way of breaking up a species to form two other species (this is a *dissociation reaction*). To understand how we enumerated these reactions, let's focus on association reactions and first take the simple case of the binding reactions between monomers. The stacked trimer structure has a total of 6 positions that could be filled with monomeric subunits (species $S_1$) and, as depicted in the main text, the monomer itself has 5 free positions. The very first reaction that can take place during assembly is the binding between two monomers, which are the reactions listed in equations (1) and (2). When one monomer binds to another monomer, there are three possible ways in which the second monomer can bind to the first; these three ways are shown in **Figure S3**. Here, the second monomer can bind in either the left of right position of "top ring" of the the first monomer forming the $K_{D,1}$ dimer species $S_3$ (**Figure S3A-B**). Alternatively, the second monomer can bind in the "opposite ring" across from the first monomer forming the $K_{D,2}$ dimer species $S_2$ (**Figure S3C**). The second monomer cannot bind at any other position, as a bond would not form between the two monomers. In other words, that reaction would not form another chemical species in the list in **Figure 1C**, and therefore is not a valid association reaction.

Note that there are two possibilities for forming the $K_{D,1}$ bond: the monomer can bind to either the left or to the right. Regardless of where it binds, however, this always forms exactly the same chemical species. This means there are two "ways" that this reaction can go forward, compared to only one way for the reaction in **Figure S3C**. We call this the "combinatorial coefficient" $c$ of the reaction, and it represents the number of ways in which two reactants can combine to form *the same* chemical species. As such, we group the two reactions in **Figure S3A-B** as a single reaction since they both result in the same product, species $S_3$. Enumerating all association reactions is relatively simple; we take every possible pair of chemical species in **Figure 1C** from the main text and determine if they can react and, if so, how many different ways in which they can react.

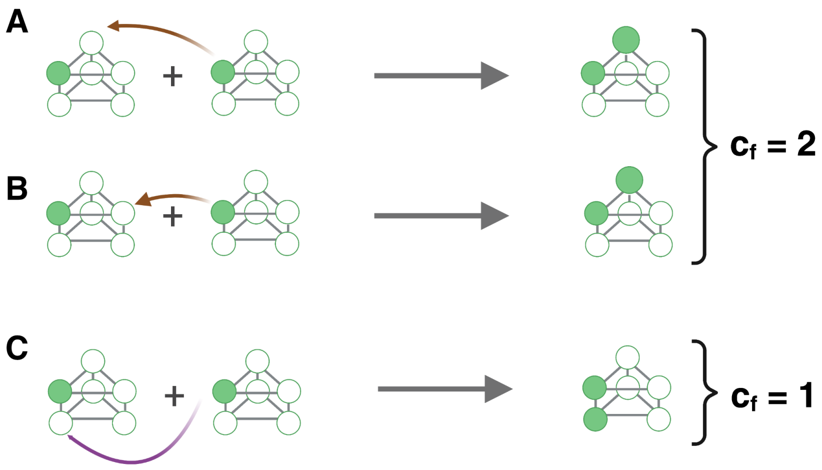

**Figure S3: Monomer binding and combinatorial coefficient.** Chemical reactions when binding two monomers species $S_1$. Following the same format as the main text, the filled in circles represent a present monomer and the unfilled circles represent an available location for a new monomer. (**A**) A second monomer can bind in the left location next to the first monomer and form the $K_{D,1}$ dimer species $S_3$. (**B**) A second monomer can bin in the right location next to the first monomer and also form the $k_{d,1}$ dimer. (**C**) A second monomer can bind to the location across from the first monomer and form the $K_{D,2}$ dimer species $S_2$. (**D**) Example of a disassociation event where a trimer ring can disassemble into a dimer and monomer in three different ways that result in the same products. The combinatorial coefficient $c_f$ is included on the right hand side of reactions. Figure made with BioRender.

This involves choosing one of the pair and fixing its orientation, and then rotating and "flipping" the second species in all possible ways and seeing if it produces a valid chemical species. The reaction is only valid if there are: (A) no steric clashes in the resulting structure and (B) if the reaction forms at least one non-covalent bond. See **Figure S**4 for a few examples of reactions that would generate a clash. For convenience, we denote the number of $K_{D,1}$ bonds formed in a reaction as $n_1$ and the number of $K_{D,2}$ bonds formed as $n_2$.

Similarly, during a dissociation reaction, there can be more than one way to dismantle an intermediate. For example, consider the single ring intermediate species $S_9$. In **Figure S**5 we see that there are three different ways in which the ring can disassemble into a dimer and a monomer. This indicates that the combinatorial coefficient for this reaction is $c = 3$ **Table S1**. Note that every forward reaction has a reverse reaction and *vice versa*, so in enumerating all the forward reactions we have also enumerated all the backwards reactions. All that remains is to determine, for any given dissociation reaction, its corresponding combinatorial coefficient. This is just the number of different ways of breaking bonds in the product structure so that the two resulting species are the two reactants on the left-hand side of the reaction.

To assist in our discussion of how we developed our ODEs, it is helpful to introduce a slightly more formal approach to our discussion of the chemical reactions in this case. Any reaction described above can be represented as a tuple: $(R, P, \nu, c_f, c_b, k_f, k_b)$ where $R, P \subset S$ (recall $S$ is the set of chemical species) are the sets of reactants and products, respectively, $\nu$ is a function that maps each species in a reaction to its stoichiometric coefficient ($\nu : S \longrightarrow \mathbb{Z}_+$), $c_f$ and $c_b$ are the combinatorial coefficients of the forward and backwards reactions, and $k_f, k_b \in \mathbb{R}_+$ are the reaction rate constants for the reaction. In **Table S1**, we list of all the reactions associated with the assembly of a stacked trimer. Note that each listed reaction is a pair of both association and disassociation reactions, and the corresponding combinatorial coefficients ($c_f$ and $c_b$) are listed as well. We also include the number of $K_{D,1}$ and $K_{D,2}$ bonds formed in each reaction as $n_1$ and $n_2$, respectively.

#### 1.4 Calculating dissociation rates

As described above, the two dissociation constants that define the affinity of the two types of non-covalent bonds in the structure are defined as the ratio of these $k_f$ and $k_b$ values for the two different reactions that lead to dimer formation (**Figure S**3). As described in the main text, one of the main areas of interest of our work is in varying these $K_D$ values and determining how that influences assembly. In varying these values, we could either vary $k_f$, $k_b$, or both. In our previous work, we argued that, for any given structure, it is likely that $k_b$ is the primary variable that influences $K_D$ over the course of evolution [3]. Thus, when we vary $K_D$ in this case, we do so by varying $k_b$ while $k_f$ is fixed. In addition, it is possible that $k_f$ will vary between reactions based on the size and structure of the intermediates. Unfortunately, to our knowledge no systematic theoretical exploration of

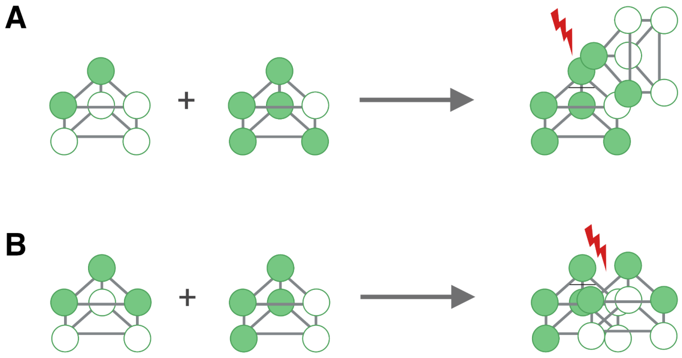

**Figure S4: Incompatible species.** Examples of reactions that cannot take place during stacked trimer assembly. **(A)** Reaction between a dimer $S_3$ and an intermediate missing only one monomer $S_{11}$. **(B)** Reaction between the top ring $S_9$ and a tetramer $S_8$. The red bolt signifies that these intermediates cannot form a new structure as there is steric clash between the structures. Figure made with BioRender.

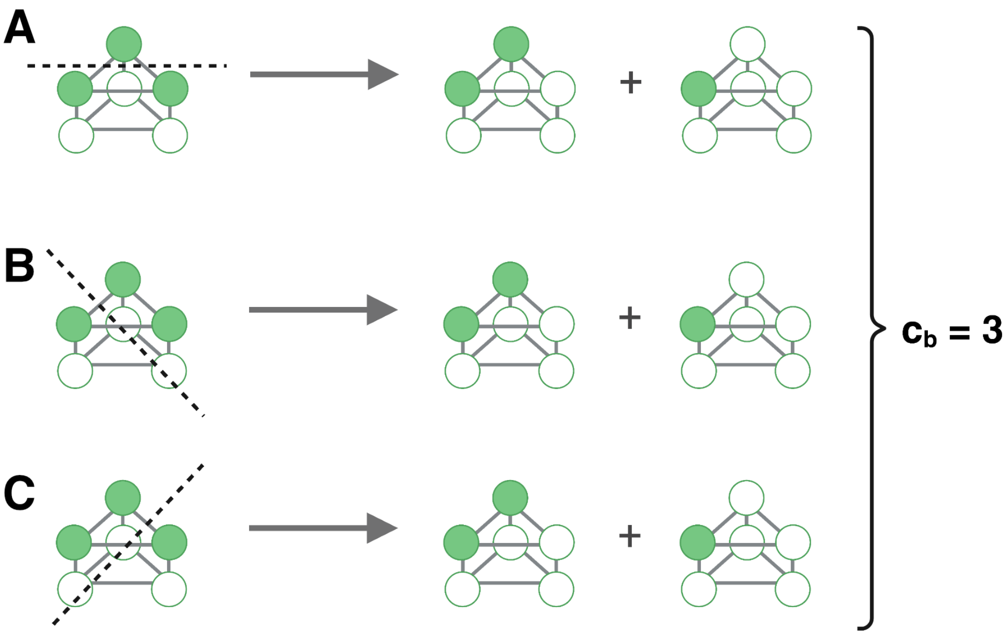

**Figure S5: Intermediate unbinding.** Chemical reactions when an intermediate disassociates. In this example, a trimeric ring, Species $S_9$ can disassemble into a dimer $S_3$ and monomer $S_1$ in three different ways that result in the same products. The dashed line highlights the bonds that break for a single monomer to unbind from the ring. One of the three monomers can unbind from the ring from the **(A)** top, **(B)** left or **(C)** right. The combinatorial coefficient $c_b$ is included for this set of reactions. Figure made with BioRender.

| Table S1: All reactions and their tuples | | | | |
|---|---|---|---|---|
| **Reaction** | $c_f$ | $c_b$ | $K_{d,1}$ | $K_{d,2}$ |
| 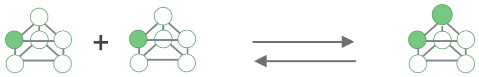 | 2 | 1 | 1 | 0 |
| 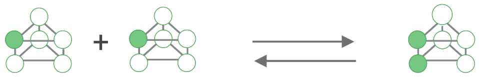 | 1 | 1 | 0 | 1 |
| 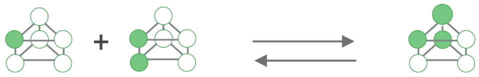 | 2 | 1 | 1 | 0 |
| 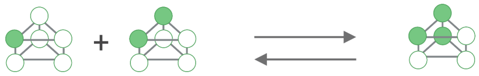 | 1 | 1 | 0 | 1 |
| 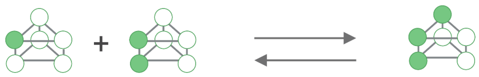 | 2 | 1 | 1 | 0 |
| 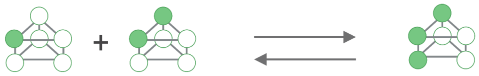 | 1 | 1 | 0 | 1 |
| 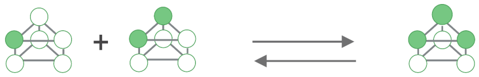 | 1 | 3 | 2 | 0 |
| 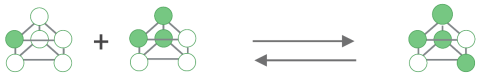 | 1 | 2 | 1 | 0 |
| 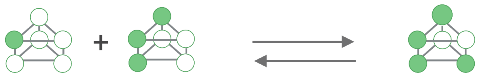 | 1 | 2 | 1 | 0 |
| 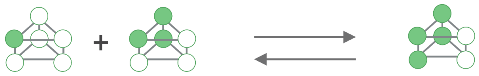 | 1 | 2 | 1 | 1 |
| 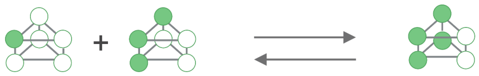 | 1 | 2 | 1 | 1 |

| Table S1: All reactions and their tuples | | | | |
|---|---|---|---|---|
| **Reaction** | $c_f$ | $c_b$ | $K_{d,1}$ | $K_{d,2}$ |
| 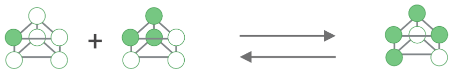 | 1 | 1 | 2 | 0 |
| 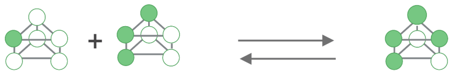 | 1 | 1 | 2 | 0 |
| 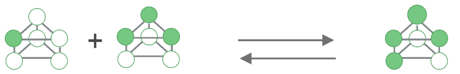 | 3 | 1 | 0 | 1 |
| 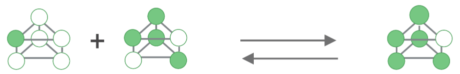 | 2 | 1 | 2 | 1 |
| 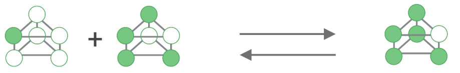 | 2 | 1 | 2 | 1 |
| 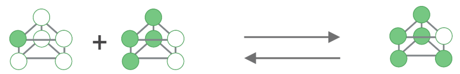 | 2 | 1 | 2 | 0 |
| 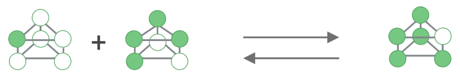 | 2 | 2 | 1 | 1 |
| 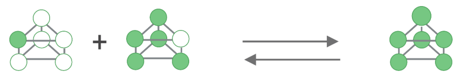 | 1 | 6 | 2 | 1 |
| 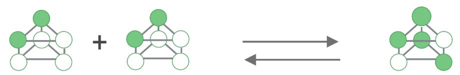 | 1 | 1 | 0 | 1 |
| 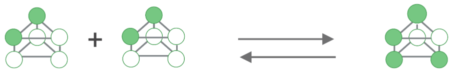 | 1 | 1 | 0 | 1 |
| 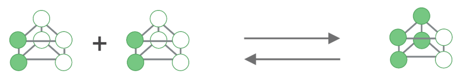 | 4 | 1 | 2 | 0 |
| 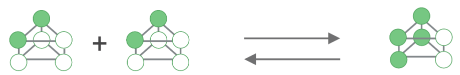 | 1 | 1 | 0 | 2 |

**Table S1: All reactions and their tuples**

| Reaction | $c_f$ | $c_b$ | $K_{d,1}$ | $K_{d,2}$ |
|---|---|---|---|---|
| 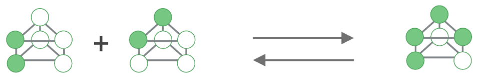 | 2 | 1 | 2 | 0 |
| 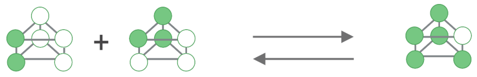 | 2 | 1 | 3 | 0 |
| 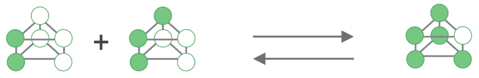 | 2 | 1 | 3 | 0 |
| 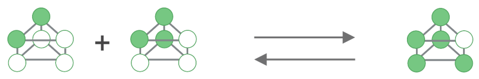 | 1 | 1 | 2 | 1 |
| 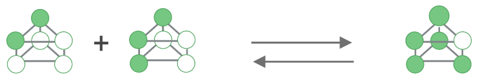 | 1 | 1 | 2 | 1 |
| 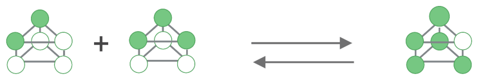 | 3 | 1 | 0 | 2 |
| 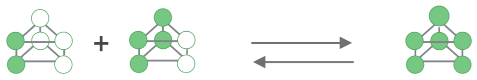 | 2 | 3 | 4 | 0 |
| 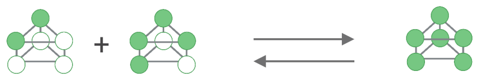 | 1 | 6 | 2 | 2 |
| 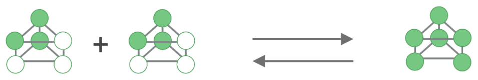 | 1 | 3 | 4 | 1 |
| 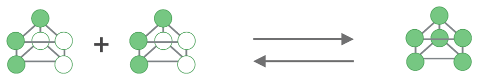 | 1 | 3 | 4 | 1 |
| 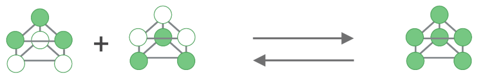 | 3 | 1 | 0 | 3 |

the influence of size and structure on effective association rates in macromolecular assembly has ever been done. As such, it is difficult to predict the relative values of $k_f$ for each possible reaction.

Even though the value of $k_f$ could theoretically vary between different reactions, since the actual interfaces are the same for each reaction, they would not have different electrostatic properties, and thus the variation in $k_f$ is likely much lower than the orders of magnitude variation in $k_b$ explored here. As such, we take $k_f$ to be a constant for all the reactions in the system [3, 14, 17]. In contrast, however, some of the dissociation reactions involve the disruption of more than one interface. For instance, the dissociation of a monomer from a single ring involves breaking two distinct $K_{D,1}$ bonds (**Figure S5**). Extensive thermodynamic calculations have shown that ring-like structures are extremely stable, so dissociation of a monomer from a single ring will occur on much longer timescales than dissociation of a dimer [3, 14] . The values of $k_b$ thus varies significantly for different reactions.

To calculate the value of $k_b$, we generalized the thermodynamic calculations for single rings in previous work [3, 14]. To explain that generalization, we first start with the simplest case, a reaction that forms a single interface. Take Reaction (1) where two monomers form the $K_{D,1}$ dimer with the formation of a single bond ($n_1 = 1$ and $n_2 = 0$). The standard free energy of formation for the $K_{D,1}$ bond type is $\Delta G_1^0$, which contains both enthalpic and entropic contributions. In this case, it is helpful to decompose the standard free energy of this reaction into two distinct parts:

$$\Delta G_1^0 = \Delta G_{I,1}^0 + \Delta G_{P,1}^0,$$

where $\Delta G_{I,1}^0$ is the free energy of formation of the interaction interface and $\Delta G_{P,1}^0$ is the loss of positional entropy [3, 14]. The former captures all intermolecular interactions between the two molecules, as well as the desolvation of the relevant interfaces, and of course for this reaction to proceed this will generally be a negative term. The positional entropy refers to the fact that, before the reaction occurs, the two monomers can adopt any relative orientation. When they are bound, however, knowledge of the position of one of the monomers allows one to predict the position of the second monomer with high accuracy. This is one of the main thermodynamic forces opposing the binding reaction, and so $\Delta G_P^0$ is positive. Estimates for the values of $\Delta G_P^0$ vary, but MD calculations for proteins suggest a value of around 9 kcal mol$^{-1}$ [3, 8, 14]. In this work, we assume the rigidity of all intermediates is roughly the same, and so we use the same value of $\Delta G_P$ for all reactions. As such, we remove the subscript here and use a constant $\Delta G_P = 9$ kcal mol$^{-1}$ for all reactions.

The dissociation constant for this reaction, $K_{D,1}$, can be calculated based on the free energy of formation of the bond, $K_{D,1} = c_0 e^{\Delta G_1^0/RT}$, where $c_0 = 1$ M and $RT$ is the product of the molar gas constant and temperature. Note that, in this work, we set the values of $K_{D,1}$ and $K_{D,2}$ and determine how this impacts assembly dynamics and efficiency (e.g. **Figure 2** in the main text). Since $k_f$ is considered a constant, this allows us to calculate $k_b$ for both the reactions (e.g. $k_b$ for Reaction 1 is simply $k_b = k_f K_{D,1}$). Note that this also corresponds to setting values for $\Delta G_1^0$ and $\Delta G_2^0$ and the free energy of formation of the corresponding interfaces.

The question then becomes how to handle reactions where more than one bond of any given type is formed. To explain how to calculate dissociation rates in this case, first consider the disruption of two $K_{D,1}$ bonds (**Figure S5**). For this reaction, $n_1 = 2$ and $n_2 = 0$. Note that, in this case, the forward reaction involves forming two $K_{D,1}$ interfaces, both of which will contribute to the free energy of the corresponding reaction. The reaction only involves the loss of the positional entropy of a single monomer, however; as a result, this reaction involves the formation of two interfaces for the price of constraining the movement of only a single monomer. As such, we can write the standard free energy of this reaction as:

$$\begin{aligned} \Delta G^0 &= 2\Delta G_{I,1}^0 + \Delta G_P^0 \\ &= 2\Delta G_1^0 - \Delta G_P, \end{aligned}$$

where the second equation follows in a straightforward way from the decomposition of $\Delta G_1^0$.The dissociation constant $K_D$ for this reaction is given by:

$$\begin{aligned} K_D &= c_0 e^{\Delta G^0/RT} \\ &= c_0 e^{\frac{2\Delta G_1^0 - \Delta G_P}{RT}} \\ &= c_0 e^{\left(\frac{2\Delta G_1^0}{RT}\right)} e^{\left(\frac{-\Delta G_P}{RT}\right)} \\ &= (K_{D,1})^2 K_P, \end{aligned}$$

where $K_P$ is a constant defined as $K_P \equiv e^{\left(\frac{-\Delta G_P}{RT}\right)}$. This calculation immediately suggests that ring-like structures are extremely thermodynamically stable. Assume $RT = 0.6$ kcal mol$^{-1}$ at room temperature and $\Delta G_P = 9$ kcal mol$^{-1}$. If $K_{D,1} = 10^{-6}$ M, then the dissociation constant for the reaction in **Figure S5** is $\sim 3 \times 10^{-19}$ M.

The above derivation is identical to that described in previous work for single rings [3, 14]. In our system, more than two bonds can be broken at a time, and these bonds can be of different types. The above logic, however, generalizes in an extremely straightforward way to this more complex case. Consider a reaction that forms $n_1$ $K_{D,1}$ bonds and $n_2$ $K_{D,2}$ bonds. Then we can write the standard free energy of this reaction as:

$$\Delta G^0(n_1, n_2) = n_1 \Delta G^0_{I,1} + n_2 \Delta G^0_{I,2} + \Delta G_P.$$

Note that, as in the above case, we are forming multiple interfaces (now of multiple types), but losing the positional entropy of only one molecule. We can re-write the above equation using just the free energies of association of the two bonds:

$$\Delta G^0(n_1, n_2) = n_1 \Delta G^0_1 + n_2 \Delta G^0_2 - (n_1 + n_2 - 1)\Delta G_P,$$

where the last term ensures that we are only losing 1 "unit" of positional entropy. We can write an effective dissociation constant for this reaction as:

$$K_{eff}(n_1, n_2) = (K_{D,1})^{n_1} (K_{D,2})^{n_2} K_P^{n_1+n_2-1}. \qquad (5)$$

For any given reaction that forms $n_1$ and $n_2$ bonds of the corresponding types, we can then calculate $k_b$ for that reaction as $k_b \equiv k_e f f(n_1, n_2) = k_f K_{eff}(n_1, n_2)$.

While the above derivation is straightforward, we should highlight that we are making several critical assumptions. For one, holding $k_f$ constant for all reactions dramatically simplifies the problem, but we may miss important dynamic features that arise from differences in association rates between reactions. We leave exploration of these differences, which will almost certainly involve detailed biophysical simulations, to future work [1, 11, 12, 13, 16, 15]. The thermodynamic calculation used here essentially assumes that the various intermediate structures are fairly rigid, and neglects differences in conformational fluctuations that could occur between various intermediates. Relaxing this assumption could again lead to interesting differences in assembly dynamics. That being said, the results described here are relatively insensitive to the specific value of $\Delta G_P$ that we choose. For instance, if $K_{D,1} = 10^{-6}$ M and $\Delta G_P = 9$ kcal mol$^{-1}$, then $K_{eff}(2, 0) \sim 3 \times 10^{-19}$ M as described above. If we dramatically reduce $\Delta G_P$ to, say, 6 kcal mol$^{-1}$, then we have $K_{eff}(2, 0) \sim 5 \times 10^{-17}$ M. While this is a change of two orders of magnitude, this intermediate is still so stable that this difference has no impact on assembly dynamics occurring on biologically-relevant timescales. As such, the specific values of the positional entropy loss we assume have little impact on our results.

#### 1.5 Ordinary Differential Equations for Stacked Trimer Assembly in the *in vitro* and *in vivo* models

In the sections above, we described how we enumerated all the possible chemical species that can occur in the assembly of the stacked trimer, as well as all the reactions that could occur between them. Call this set of species $S$ and the set of reactions $\mathcal{R}$ (see **Figure 1** and **Table S1**, respectively). Let $x_j$ be the concentration of species $S_j$ at some particular time $t$. Our goal here is to determine $x'_j$, or the change in $x_j$ over time, given our set of reactions $\mathcal{R}$. As described above, each reaction $r_i \in \mathcal{R}$ is a tuple defined as:

$$r_i = (R, P, \nu, c_f, c_b, k_f, n_1, n_2)$$

where $R$ is the set of reactants and $P$ is the set of products for the reaction. Here, $\nu$ is the stoichiometric function as described above that maps any given species in the set of reactants or products to the stoichiometric coefficient it has in the reaction (i.e. the number of molecules of that species consumed or produced by that reaction, $\nu : R \cup P \longrightarrow \mathbb{Z}_+$). As an example, consider Reaction (1) where $r_1 = (\{S_1\}, \{S_3\}, \nu, 1, 1, k_f, k_b)$ and

$$\nu(S_1) = 2$$
$$\nu(S_3) = 1.$$

Every reaction in $\mathcal{R}$ has this structure. To see how $x_j$ changes over time, we need to identify the reactions that involve $x_j$. To simplify the definition of the ODEs, we will follow the convention that the forward reaction is the binding reaction and the backward reaction is the unbinding reaction. As a result, for all the reactions in $\mathcal{R}$, the left-hand side contains smaller intermediates and the right-hand side contains the larger products resulting from the binding reaction. Let $\phi(S_j)$ be the set of reactions where $S_j$ is the product and $\theta(S_j)$ be the set of reactions where $S_j$ is a reactant, so:

$$\phi(S_j) = \{r_i | S_j \in P_i\}$$
$$\theta(S_j) = \{r_i | S_j \in R_i\},$$

where every $r_i \in \mathcal{R}$ and the sets $P_i$ and $R_i$ should be clear. Note that, given the above definition, we have $\phi : S \rightarrow \mathcal{P}(\mathcal{R})$ and $\theta : S \rightarrow \mathcal{P}(\mathcal{R})$, where $\mathcal{P}(X)$ is the power set of some set $X$. In other words, these two functions map each species to the subset of reactions where that species is a product or reactant, respectively. To illustrate these functions, imagine a simple scenario where the only reaction possible in the system was Reaction (1), i.e. $\mathcal{R} = \{r_1\}$. Here we can see that $\phi(S_1) = \emptyset$ and $\phi(S_2) = \{r_1\}$. Similarly, $\theta(S_1) = \{r_1\}$ and $\theta(S_2) = \emptyset$.

Given these definitions, we can now turn to writing $x'_j$. The first component of our equation for this particular ODE will involve all the reactions where the species corresponding to this concentration, $S_j$, is a *product* of that reaction. So this will just be the set of reactions returned by the function $\phi(S_j)$. To simplify writing the ODEs, we introduce a final definition, which we call the "Fundamental Fluxes" of a reaction; each reaction $r_i$ has a fundamental forward flux, $\mathcal{F}_{f,i}$, and a fundamental reverse flux $\mathcal{F}_{b,i}$. Using the law of mass action associated with species $S_i$ production, $\mathcal{F}_{f,i}$ is given by:

$$\mathcal{F}_{f,i} = k_f c_f \prod_{S_k \in R_i} \frac{x_k^{\nu_i(S_k)}}{\nu_i(S_k)!}.$$

Note that this presentation may be slightly different from the standard approach to defining the law of mass action. The reason for this has to do with the fact that choosing a single, constant $k_f$ for all reactions requires that attention be paid to combinatorial factors, which leads, for instance, to the factorial term within the product. A detailed explanation for how these terms derive from the stochastic law of mass action may be found in the supplement to ref. [3]. For the example with $r_1$, we can calculate the forward fundamental flux as follows:

$$\begin{aligned}\mathcal{F}_{f,1} &= k_f c_f \prod_{S_k \in R_1} \frac{x_k^{\nu(S_k)}}{\nu(S_k)!} \\ &= k_f * 1 * \frac{x_1^2}{2!} \\ &= \frac{1}{2} k_f x_1^2\end{aligned}$$

Similarly,the reverse fundamental flux $\mathcal{F}_{b,i}$ is given by:

$$\mathcal{F}_{b,i} = k_{eff}(n_1, n_2) c_b \prod_{S_k \in P_i} \frac{x_k^{\nu_i(S_k)}}{\nu_i(S_k)!}.$$

Recall that $n_1$ and $n_2$ represent the number of $K_{D,1}$ and $K_{D,2}$ bonds are formed in a reaction, and we calculate the effective dissociation rate $k_{eff}$ using those numbers (see section 1.4 above). To continue with the example with Reaction (1), the reverse fundamental flux can be calculated as:

$$\begin{aligned}\mathcal{F}_{b,1} &= k_{eff}(n_1, n_2) c_b \prod_{S_k \in P_1} \frac{x_k^{\nu(S_k)}}{\nu(S_k)!} \\ &= k_{eff}(0,1) * 1 * \frac{x_2^1}{1!} \\ &= k_{eff}(0,1) x_2.\end{aligned}$$

Again, for a more detailed description of the $\nu_i(S_k)!$ term, refer to the discussion in the supplement of ref. [3].

Finally, we can define $x'_j$ as the product of the fluxes and the stoichiometric coefficients as:

$$x'_j = \sum_{r_i \in \phi(S_j)} \nu_i(S_k) \left(\mathcal{F}_{f,i} - \mathcal{F}_{b,i}\right) + \sum_{r_i \in \theta(S_j)} \nu_i(S_j) \left(-\mathcal{F}_{f,i} + \mathcal{F}_{b,i}\right). \tag{6}$$

Note that the first sum in this case sums over all the reactions where $S_j$ is the product of the reaction, and the second sum considers all the cases where $S_j$ is a reactant. The two fluxes in each case arise due to the fact that every reaction is *reversible*; in other words, when the species is a product, its concentration is increased by the forward (binding) reaction and decreased by the reverse (unbinding) reaction. The opposite holds for cases where the species in question is a reactant. To make this more clear, consider again the case where Reaction (1) is the only reaction that can occur. We can use Equation 6 to write the ODEs for both $x_1$ and $x_2$ as:

$$
\begin{aligned}
x_1' &= \sum_{r_i \in \phi(S_1)} \left(\mathcal{F}_{f,i} - \mathcal{F}_{b,i}\right) + \sum_{r_i \in \theta(S_1)} \left(-\mathcal{F}_{f,i} + \mathcal{F}_{b,i}\right) \\
&= 0 + 2\left(-\frac{1}{2}k_f x_1^2 + k_{eff}(0,1)x_2\right) \\
&= -k_f x_1^2 + 2k_{eff}(0,1)x_2 \\
x_2' &= \sum_{r_i \in \phi(S_1)} \left(\mathcal{F}_{f,i} - \mathcal{F}_{b,i}\right) + \sum_{r_i \in \theta(S_1)} \left(-\mathcal{F}_{f,i} + \mathcal{F}_{b,i}\right) \\
&= \frac{1}{2}k_f x_1^2 - k_{eff}(0,1)x_2 + 0
\end{aligned}
$$

Notice that $x_1' = -2x_2'$, which directly reflects the condition for the conservation of mass. Fore more details on how the ODEs are generated, refer to [3]. The list of ODEs for the stacked trimer model is below:

$$
\begin{aligned}
x_1' = &-2k_f * 0.5 * x_1x_1 + 1k_{eff}(1,0)x_2 - 2k_f * 0.5 * x_1x_1 + 1k_{eff}(1,0)x_2 - 1k_f * 0.5 * x_1x_1 \\
&+ 1k_{eff}(0,1)x_3 - 1k_f * 0.5 * x_1x_1 + 1k_{eff}(0,1)x_3 - 1k_f x_1 * x_2 + 3k_{eff}(2,0)x_4 - 1k_f x_1 * x_2 \\
&+ 1k_{eff}(0,1)x_5 - 1k_f x_1 * x_2 + 1k_{eff}(0,1)x_6 - 2k_f x_1 * x_3 + 1k_{eff}(1,0)x_6 - 2k_f x_1 * x_3 \\
&+ 1k_{eff}(1,0)x_5 - 3k_f x_1 * x_4 + 1k_{eff}(0,1)x_7 - 1k_f x_1 * x_5 + 1k_{eff}(2,0)x_7 - 1k_f x_1 * x_5 \\
&+ 2k_{eff}(1,0)x_8 - 1k_f x_1 * x_5 + 2k_{eff}(1,1)x_9 - 1k_f x_1 * x_6 + 1k_{eff}(2,0)x_7 - 1k_f x_1 * x_6 \\
&+ 2k_{eff}(1,1)x_9 - 1k_f x_1 * x_6 + 2k_{eff}(1,0)x_{10} - 2k_f x_1 * x_7 + 2k_{eff}(1,1)x_{11} - 2k_f x_1 * x_8 \\
&+ 1k_{eff}(2,1)x_{11} - 2k_f x_1 * x_9 + 1k_{eff}(2,0)x_{11} - 2k_f x_1 * x_{10} + 1k_{eff}(2,1)x_{11} - 1k_f x_1 * x_{11} \\
&+ 6k_{eff}(2,1)x_{12} \\
x_2' = &\ 2k_f * 0.5 * x_1x_1 - 1k_{eff}(1,0)x_2 - 1k_f x_1 * x_2 + 3k_{eff}(2,0)x_4 - 1k_f x_1 * x_2 \\
&+ 1k_{eff}(0,1)x_5 - 1k_f x_1 * x_2 + 1k_{eff}(0,1)x_6 - 1k_f * 0.5 * x_2 * x_2 + 1k_{eff}(0,2)x_9 \\
&- 1k_f * 0.5 * x_2 * x_2 + 1k_{eff}(0,2)x_9 - 1k_f * 0.5 * x_2 * x_2 + 1k_{eff}(0,1)x_{10} - 1k_f * 0.5 * x_2 * x_2 \\
&+ 1k_{eff}(0,1)x_{10} - 1k_f * 0.5 * x_2 * x_2 + 1k_{eff}(0,1)x_8 - 1k_f * 0.5 * x_2 * x_2 + 1k_{eff}(0,1)x_8 \\
&- 2k_f x_2 * x_3 + 1k_{eff}(2,0)x_7 - 3k_f x_2 * x_4 + 1k_{eff}(0,2)x_{11} - 1k_f x_2 * x_5 + 1k_{eff}(2,1)x_{11} \\
&- 1k_f x_2 * x_6 + 1k_{eff}(2,1)x_{11} - 1k_f x_2 * x_7 + 6k_{eff}(2,2)x_{12} \\
x_3' = &\ 1k_f * 0.5 * x_1x_1 - 1k_{eff}(0,1)x_3 - 2k_f x_1 * x_3 + 1k_{eff}(1,0)x_6 - 2k_f x_1 * x_3 + 1k_{eff}(1,0)x_5 \\
&- 2k_f x_2 * x_3 + 1k_{eff}(2,0)x_7 - 4k_f * 0.5 * x_3 * x_3 + 1k_{eff}(2,0)x_9 - 4k_f * 0.5 * x_3 * x_3 \\
&+ 1k_{eff}(2,0)x_9 - 2k_f x_3 * x_5 + 1k_{eff}(3,0)x_{11} - 2k_f x_3 * x_6 + 1k_{eff}(3,0)x_{11} \\
&- 2k_f x_3 * x_9 + 3k_{eff}(4,0)x_{12} \\
x_4' = &\ 1k_f x_1 * x_2 - 3k_{eff}(2,0)x_4 - 3k_f x_1 * x_4 + 1k_{eff}(0,1)x_7 - 3k_f x_2 * x_4 \\
&+ 1k_{eff}(0,2)x_{11} - 3k_f * 0.5 * x_4 * x_4 + 1k_{eff}(0,3)x_{12} - 3k_f * 0.5 * x_4 * x_4 + 1k_{eff}(0,3)x_{12} \\
x_5' = &\ 1k_f x_1 * x_2 - 1k_{eff}(0,1)x_5 + 2k_f x_1 * x_3 - 1k_{eff}(1,0)x_5 - 1k_f x_1 * x_5 + 1k_{eff}(2,0)x_7 - \\
&1k_f x_1 * x_5 + 2k_{eff}(1,0)x_8 - 1k_f x_1 * x_5 + 2k_{eff}(1,1)x_9 - 1k_f x_2 * x_5 + 1k_{eff}(2,1)x_{11} \\
&- 2k_f x_3 * x_5 + 1k_{eff}(3,0)x_{11} - 1k_f * 0.5 * x_5 * x_5 + 3k_{eff}(4,1)x_{12} - 1k_f * 0.5 * x_5 * x_5 \\
&+ 3k_{eff}(4,1)x_{12} \\
x_6' = &\ 1k_f x_1 * x_2 - 1k_{eff}(0,1)x_6 + 2k_f x_1 * x_3 - 1k_{eff}(1,0)x_6 - 1k_f x_1 * x_6 + 1k_{eff}(2,0)x_7 \\
&- 1k_f x_1 * x_6 + 2k_{eff}(1,1)x_9 - 1k_f x_1 * x_6 + 2k_{eff}(1,0)x_{10} - 1k_f x_2 * x_6 + 1k_{eff}(2,1)x_{11} \\
&- 2k_f x_3 * x_6 + 1k_{eff}(3,0)x_{11} - 1k_f * 0.5 * x_6 * x_6 + 3k_{eff}(4,1)x_{12} - 1k_f * 0.5 * x_6 * x_6 \\
&+ 3k_{eff}(4,1)x_{12}
\end{aligned}
$$

$$
\begin{aligned}
x_7' &= 3k_f x_1 * x_4 - 1k_{eff}(0,1)x_7 + 1k_f x_1 * x_5 - 1k_{eff}(2,0)x_7 + 1k_f x_1 * x_6 - 1k_{eff}(2,0)x_7 \\
&\quad - 2k_f x_1 * x_7 + 2k_{eff}(1,1)x_{11} + 2k_f x_2 * x_3 - 1k_{eff}(2,0)x_7 - 1k_f x_2 * x_7 + 6k_{eff}(2,2)x_{12} \\
x_8' &= 1k_f x_1 * x_5 - 2k_{eff}(1,0)x_8 - 2k_f x_1 * x_8 + 1k_{eff}(2,1)x_{11} + 1k_f * 0.5 * x_2 * x_2 - 1k_{eff}(0,1)x_8 \\
x_9' &= 1k_f x_1 * x_5 - 2k_{eff}(1,1)x_9 + 1k_f x_1 * x_6 - 2k_{eff}(1,1)x_9 - 2k_f x_1 * x_9 + 1k_{eff}(2,0)x_{11} \\
&\quad + 1k_f * 0.5 * x_2 * x_2 - 1k_{eff}(0,2)x_9 + 4k_f * 0.5 * x_3 * x_3 - 1k_{eff}(2,0)x_9 - 2k_f x_3 * x_9 \\
&\quad + 3k_{eff}(4,0)x_{12} \\
x_{10}' &= 1k_f x_1 * x_6 - 2k_{eff}(1,0)x_{10} - 2k_f x_1 * x_{10} + 1k_{eff}(2,1)x_{11} + 1k_f * 0.5 * x_2 * x_2 \\
&\quad - 1k_{eff}(0,1)x_{10} \\
x_{11}' &= 2k_f x_1 * x_7 - 2k_{eff}(1,1)x_{11} + 2k_f x_1 * x_8 - 1k_{eff}(2,1)x_{11} + 2k_f x_1 * x_9 - 1k_{eff}(2,0)x_{11} \\
&\quad + 2k_f x_1 * x_{10} - 1k_{eff}(2,1)x_{11} - 1k_f x_1 * x_{11} + 6k_{eff}(2,1)x_{12} + 3k_f x_2 * x_4 - 1k_{eff}(0,2)x_{11} \\
&\quad + 1k_f x_2 * x_5 - 1k_{eff}(2,1)x_{11} + 1k_f x_2 * x_6 - 1k_{eff}(2,1)x_{11} + 2k_f x_3 * x_5 - 1k_{eff}(3,0)x_{11} \\
&\quad + 2k_f x_3 * x_6 - 1k_{eff}(3,0)x_{11} \\
x_{12}' &= 1k_f x_1 * x_{11} - 6k_{eff}(2,1)x_{12} + 1k_f x_2 * x_7 - 6k_{eff}(2,2)x_{12} + 2k_f x_3 * x_9 - 3k_{eff}(4,0)x_{12} \\
&\quad + 3k_f * 0.5 * x_4 * x_4 - 1k_{eff}(0,3)x_{12} + 1k_f * 0.5 * x_5 * x_5 - 3k_{eff}(4,1)x_{12} + 1k_f * 0.5 * x_6 * x_6 \\
&\quad - 3k_{eff}(4,1)x_{12}
\end{aligned}
$$

The list of ODEs above represents the "*in vitro*" model discussed in the main text. We added synthesis of monomers and degradation of all the chemical species to generate the "*in vivo*" model. In this model, we assume that the monomers (e.g. Species $S_1$) is produced by the cell at a constant rate, and we call this $0^{\text{th}}$-order rate constant $Q$. Since cells clearly cannot directly synthesize protein complexes, this assumption of monomer synthesis is quite natural. Degradation, however, can take several different forms. In this model, we consider the simplest case where all chemical species are degraded at the same rate, regardless of size or structure. This models the case where loss of proteins occurs mostly through dilution due to cell growth [3, 6, 10], or a case where protein complexes are degraded actively by the cell (through the ubiquitin-proteasome system) with each complex degraded at the same rate. We leave exploration of other forms of degradation to further work. We call the uniform $1^{\text{st}}$-order degradation rate $\delta$. The *in vivo* model is thus a straightforward modification of the *in vitro* model, where we add the $Q$ and $\delta$ terms to the monomer ODE and the degradation term to the equations for all the other species with index $j > 1$:

$$
\begin{aligned}
x_1' &= \sum_{r_i \in \phi(S_1)} \left(\mathcal{F}_{f,i} - \mathcal{F}_{b,i}\right) + \sum_{r_i \in \theta(S_1)} \left(-\mathcal{F}_{f,i} + \mathcal{F}_{b,i}\right) + Q - \delta x_1 \\
x_j' &= \sum_{r_i \in \phi(S_k)} \nu_i(S_k) \left(\mathcal{F}_{f,i} - \mathcal{F}_{b,i}\right) + \sum_{r_i \in \theta(S_j)} \nu_i(S_j) \left(-\mathcal{F}_{f,i} + \mathcal{F}_{b,i}\right) - \delta x_j.
\end{aligned}
$$

Since this is just a straightforward addition of terms to the ODEs listed for the *in vitro* model above, we do not provide a separate full set of differential equations here.

#### 2 THE ASSEMBLY DYNAMICS OF STACKED HOMOMERIC RINGS

As discussed in the main text, the assembly time course depends on the binding affinities, $K_{D,1}$ and $K_{D,2}$, and their relative values. Below are time course plots similar to **Figure 2** in the main text. However, **Figure** S6 includes the assembly yield for *all* of the intermediates for the *in vitro* model. Each panel in **Figure** S6 includes the time course for all the species for fixed values of $K_{D,1}$ and $K_{D,2}$.

#### 3 OBSERVED STACKED TRIMER STRUCTURES AVOID HAVING TWO STRONG INTERACTIONS

##### 3.1 Structural analysis of stacked trimers

Our models predicted that evolved structures would likely avoid having both $K_{D,1}$ and $K_{D,2}$ strong, since this leads to lower yields and more deadlock in both the *in vitro* and *in vivo* models. To test this prediction, we analyzed available solved structures of stacked three member rings.

We began by identifying stacked trimers from the Protein Data Bank in Europe Proteins, Interfaces, Structures and Assemblies (PDBePISA) database. To do this, we first ran a database search on PDBePISA that included the following

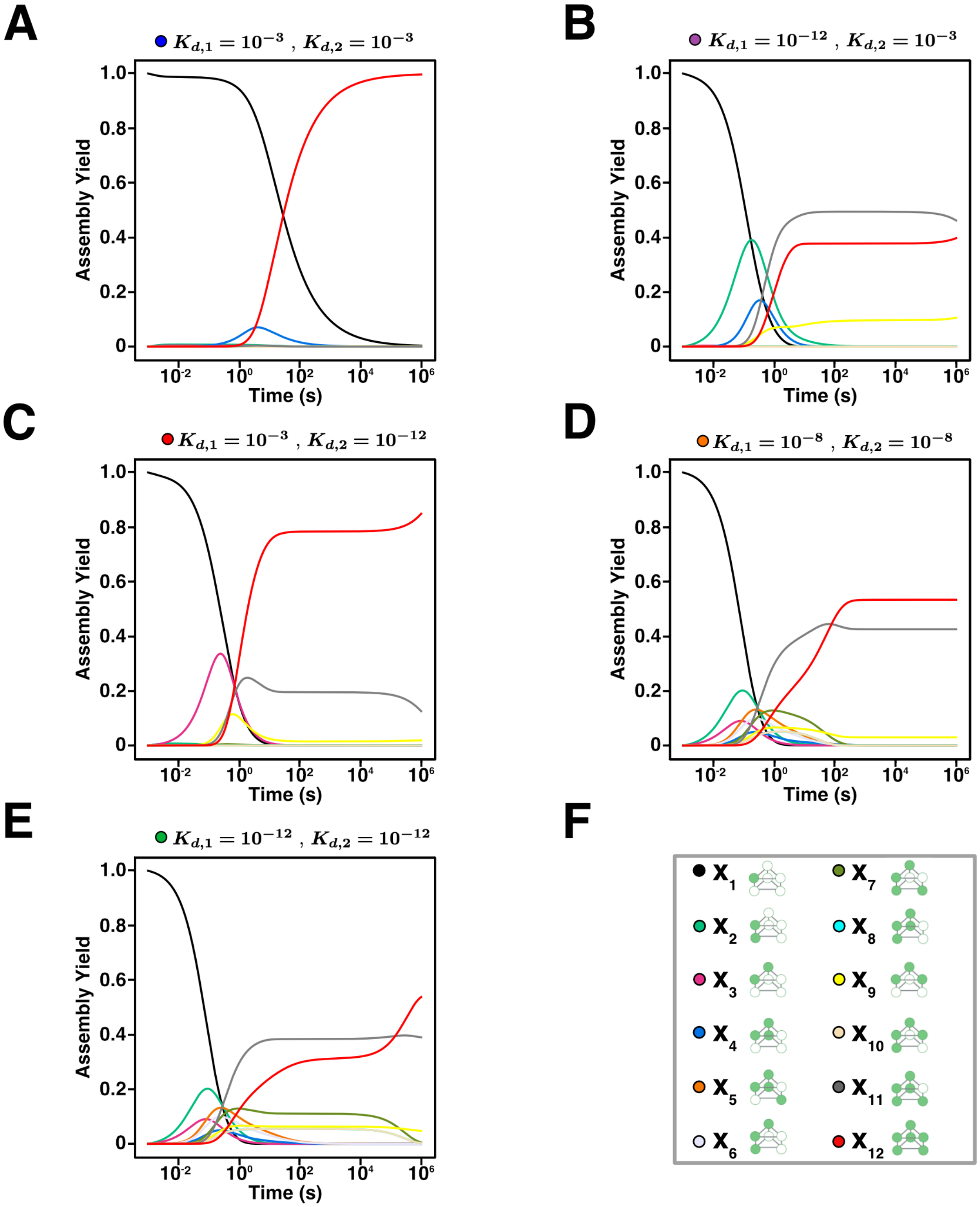

**Figure S6: Assembly time course for all intermediates in the *in vitro* model**. Each curve represents the assembly yield for each species, based on the legend in Panel **F**, over time up to $10^6$ (s) depending on fixed values of $K_{D,1}$ and $K_{D,2}$. **(A-E)** have fixed values of $K_{D,1}$ and $K_{D,2}$ labeled at the top of the plot. The colored dot at the top of the plot correspond to the assembly yield curve for the stacked trimer in the main text **Figure 2**.

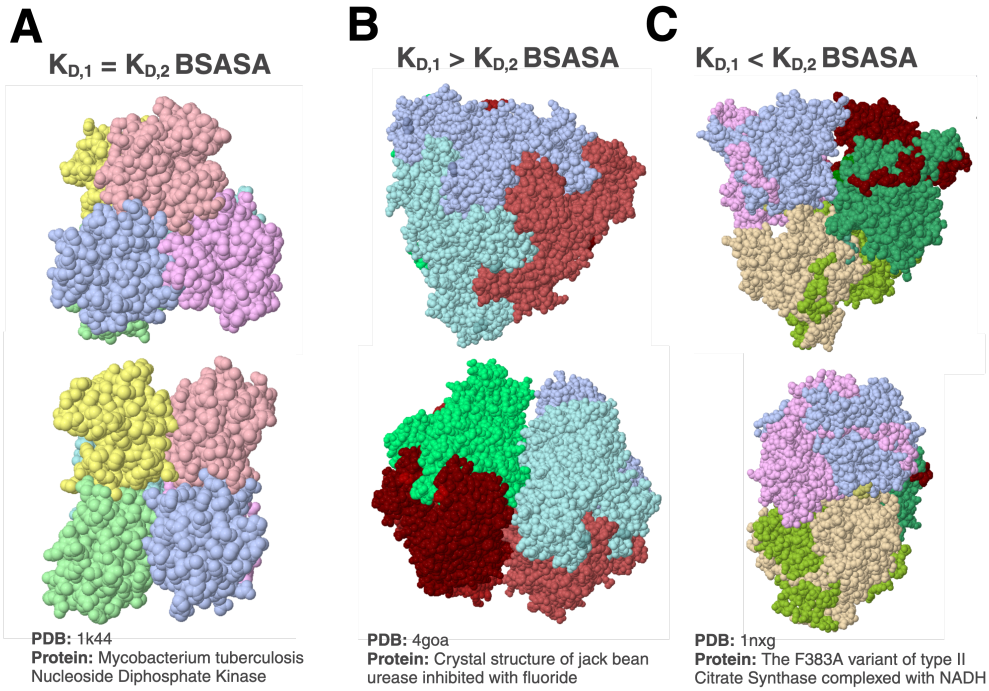

**Figure S7: Figure S6: Example structures of BSASA patterns.** Solved structures of stacked trimers. These examples include structures for which the $K_{D,1}$-BSASA is **(A)** approximately equal to the $K_{D,2}$-BSASA, **(B)** greater than the $K_{D,2}$-BSASA and **(C)** less than the $K_{D,2}$-BSASA. The protein structures that were found to have such patterns are the *Mycobacterium tuberculosis* Nucleoside Diphosphate Kinase, jack bean urease, and The F383A variant of type II Citrate Synthase. The PDB codes are included beneath each stacked trimer. Figure made with BioRender and PDBePISA.

parameters: only homomeric structures, hexamers, and only proteins as a filter. This resulted in 3619 potential stacked trimers. While this set included many structures that were *bona fide* stacked trimers, it also obviously could contain structures that were not what we were interested in, such as six-membered homomeric rings [3]. Thus, to obtain a list of only stacked trimers, we visually inspected each structure by entering each PDB entry into the PDBePISA Submission Query and analyzing it under the Assemblies option. PDBePISA provides a visual representation of the fully assembled structure; we used this feature to visually inspect the complex and determined if the topology of the complex was similar to the schematic shown in **Figure S 1a**. If a structure was determined to be a stacked trimer, we proceeded to analyze the binding affinity patterns. This resulted in a total of 1580 stacked trimers. A list of these structures with their PDB IDs is provided in Appendix SAX. While this process of visual inspection is ultimately subjective, as one can see from the list of structures obtained, this approach allowed us to narrow the set of structures to a subset that very clearly represent the type of structure we are interested in.

Similar to [3], we used the Buried Solvent-Accessible Surface Area (BSASA) to evaluate relative binding affinities, which has been shown to very roughly correlate with binding affinities in previous work [2, 5]. Specifically, we obtained the BSASA provided by PDBePISA by selecting the interfaces associated with binding within the rings and between the rings. We first identified the interfaces that make up the bond within the rings; there are six such interfaces in total (three for the top ring and three for the bottom ring). Although a stacked trimer nominally has a three-fold axis of symmetry, subtle conformational differences often lead to slight variations in the BSASA of each of these interfaces. We thus took the average BSASA for all six interfaces to represent the $K_{D,1}$-BSASA. Similarly, we took the average over the three between-ring interaction to represent the $K_{D,2}$-BSASA. The resulting BSASA were used to generate **Figure 4a** in the main text.

#### 3.2 Examples of stacked trimers

As mentioned above, we found 1580 stacked trimers from PDBePISA. As described in the main text, there were stacked trimers that had a higher BSASA in the $K_{D,1}$ interface than in the $K_{D,2}$ interface, a higher BSASA in the $K_{D,2}$ interface than in the $K_{D,1}$ interface and many that had approximately equal BSASA in both interfaces. **Figure S7** includes visual examples of each.

### 4 ANALYZING ASSEMBLY PATHWAYS

#### 4.1 Defining and enumerating assembly pathways

In the field of macromolecular assembly, it is common to use the term “assembly pathway" to describe the rough order of events that occur during assembly [7, 9]. For instance, in the assembly of the proteasome from archaea, it has long been thought that the Core Particle (CP) forms first through the assembly of a complete $\alpha_7$ ring, followed by binding of $beta$ subunits to form the $\alpha_7\beta_7$ “Half Proteasome" (HP). Two HP’s then dimerize to form the CP [4, 7, 9]. While these stories regarding assembly are common, to our knowledge there has been no attempt to make the notion of an assembly pathway mathematically formal. We hypothesized that such a formalization could assist with the analysis of our results, particularly with determining exactly *why* certain patterns of affinity lead to extensive deadlock while others do not.

Using similar notation to the above sections, we can now define the **assembly pathways** that can take place during stacked trimer formation. As described informally in the main text, an assembly pathway represents how a group of subunits go from being monomers to the fully assembled structure. Here, we define more precisely an assembly pathway as a binary tree with these three conditions: (1) there is exactly one root node which represents the fully assembled structure, in our model the stacked trimer $S_{12}$; (2) the leaf nodes represent the single subunits ($S_1$) and there are exactly as many leaf nodes as there are subunits in the structure (six in our case); and (3) for every parent node $S_k$, meaning a node with two child nodes $S_i$ and $S_j$, there exists a reaction $r_i \in \mathcal{R}$ such that the child nodes represent the reactants of reaction $r_i$ and the parent node represents the product of reaction $r_i$.

To make the above definition more formal, recall that a tree is a particular type of graph $G$, where a graph is a collection of nodes or vertices (represented by a set $V$) and edges between them (forming the set $E$), so that $G = (V, E)$. Note that we typically consider the edges in a tree to be undirected, and so every element $e \in E$ can be written as an unordered pair $(v_1, v_2)$ with $v_1, v_2 \in V$. For each tree, we take the vertices to be a finite subset of the integers, which we will denote $\mathbb{Z}_n$. Let $\Gamma_m(i)$ be a function $\Gamma_m : \mathbb{Z}_n \to S$ that takes in the index $i$ of the node on a tree $m$ and returns the species associated with the node. For example, consider the two trees in **Figure** S8 that are associated with forming the $K_{D,1}$ and $K_{D,2}$ dimers and the values for $\Gamma$ beneath each. Notice that each tree has the same leaf nodes, $S_1$, but the parent node is different, which is reflected by the values of $\Gamma$ for each tree. Therefore, although each node is unique, several nodes can map to the same species.

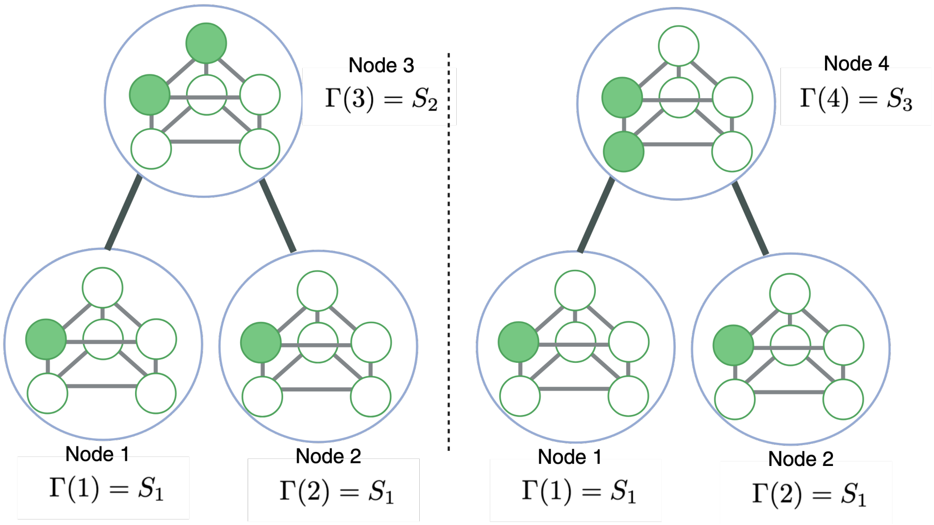

**Figure S8: Example of simple binary trees**. The trees represented here show two examples of binary trees. Suppose that both trees have the same leaf nodes but different parent nodes. Let the nodes 1 and 2 both represent the monomer species $S_1$, the node 3 represent the $K_{D,1}$ dimer and node 4 represent the $K_{D,2}$ dimer. The associated $\Gamma$ functions and evaluates are listed below each node.

An assembly pathway is particular form of graph $G$ that forms a binary tree, and there are several conditions that it needs to meet in order to be considered a valid assembly pathway. In the following discussion we will leave the terms “leaf node," “root node," “child node" and “parent node" implicit as these are standard definitions in trees. First off, we will have some unique root node; without loss of generality, assign this root node to be some element $f \in \mathbb{Z}_n$. In order for a tree $m$ to be a valid assembly pathway, we require $\Gamma_m(f) = S_{12}$; in other words, the root node represents the fully assembled structure $S_{12}$. Moreover, there are exactly six leaf nodes, and if a node $l$ is a leaf node, then $\Gamma_m(l) = S_1$, so these leaf nodes represent the

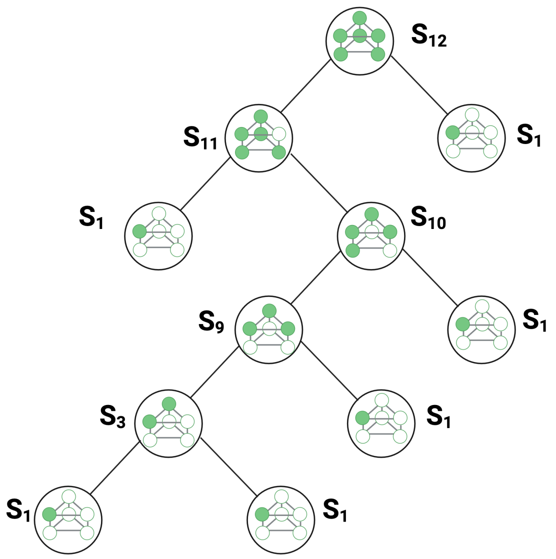

**Figure S9: Example of a pathway.** Binary tree of a pathway in which the top is the root node with the fully stacked trimer $S_{12}$. The leaf nodes at the bottom of the tree are the monomers $S_1$. The midsection of the tree are all the nodes that are assembled as intermediate structures. The root node is assembled from the intermediates $S_{11]}$ and $S_1$. Then, $S_{11}$ is assembled from $S_{12}$ and $S_1$. Intermediate $S_{10}$ is assembled from $S_9$ and $S_1$. Then, $S_9$ is assembled from $S_3$ and $S_1$. Finally, $S_3$ is assembled from two $S_1$ subunits. Note that the leaf nodes are all $S_1$ monomers are required. Figure made with BioRender.

monomers. If a node $k$ in $m$ is not a leaf node nor the root node of the tree, then by definition it will have two child nodes $i$ and $j$. In order for the tree to be an assembly pathway, there must exist some $r_q \in \mathcal{R}$ such that $\Gamma_m(i), \Gamma_m(j) \in R_q$ (i.e. the child nodes are the reactants) and $\Gamma_m(k) \in P_q$ (i.e. the parent node is the product). In other words, there has to be a reaction in the system that can *generate* the parent species from the two child species.

Intuitively, this definition of an assembly pathway makes sense: it is a set of chemical species that “starts" at the monomers (of which there are six) and “ends up" at the fully assembled structure. Every step in the pathway is a reaction that generates a larger chemical species from two smaller species, represented by the two child nodes of a given parent node. The set of all possible assembly pathways clearly represents all the scenarios whereby a stacked trimer can assemble through the chemical reactions allowed by the system. An example of a pathway is shown in **Figure** S9. Note that in this figure, as well as the pathway figures in the main text, we forgo the actual integer labels for the nodes and instead label each node with its corresponding chemical species.

Given our definition of an assembly pathway, we enumerated all of the pathways that begin with the six monomers and end with the fully assembly stacked trimer using the forward reactions listed in **Section 1.3**. We used a straightforward recursive strategy to perform this enumeration. We began by considering all of the possible reactions that can form the fully assembled stacked trimer; these represent all the possible child nodes of the root node, as seen in **Figure** S10. Then for each child node, we considered all of the reactions that can form that particular intermediate species. In other words, for each child node in **Figure** S10 we can enumerate all of its possible child nodes. We repeat this process until we reach only monomers, resulting in all possible assembly pathways. A flow chart of this process is shown in **Figure** S11. A few representative pathways are shown in the main text **Figure 5**. All 46 of the pathways that form the stacked trimer are depicted in **Figure** S12. Note that the pathways are grouped based on the final reaction in the pathway. For example, all of the pathways that have the reaction between $S_1$ and the pentamer $S_{11}$ as the last reaction in the pathway are grouped together (these constitute the last 20 pathways).

#### 4.2 Pathway contributions reveal the origins of deadlock

The ultimate goal of this analysis is to characterize how much each individual pathway *contributes* to the assembly of the stacked trimer. To do that, we start from our definition of Fundamental Fluxes in Section 1.5. As described above, any pathway $P_k$ is a binary tree, and associated with that tree are the set of forward reactions necessary for the assembly of the fully assembled structure under that scenario. Each reaction $r_i$ in that set has a fundamental flux $\mathcal{F}_{f,i}$.

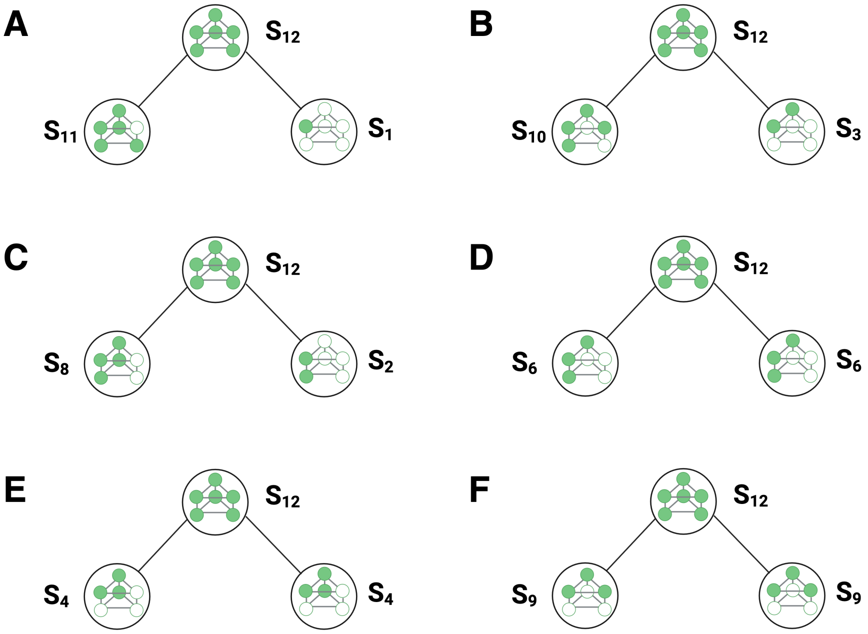

**Figure S10: Trees for reactions that form the fully stacked trimer.** Trees showing the top part of a pathway in which the fully assembled stacked trimer is formed. Each panel shows a unique step for forming the stacked trimer. For instance, in panel **(A)**, the stacked trimer $S_{12}$ can be formed from binding $S_{11}$ and $S_1$. **(B)** The stacked trimer can also be formed from binding $S_{10}$ and $S_3$ and so forth. Figure made with BioRender.

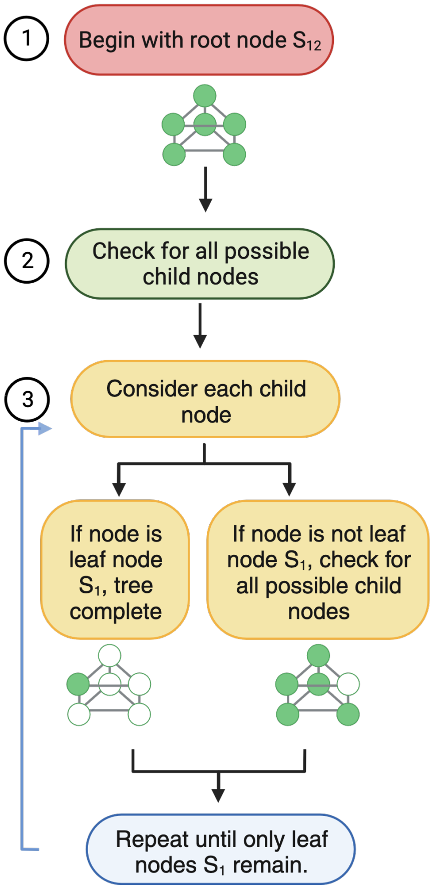

**Figure S11: Flowchart of assembly pathway enumeration.** This flowchart shows the general process to express each pathway. Begin with the root node, in the stacked trimer case, species $S_{12}$. Then check for all the possible child nodes of the root node. For each possible child node, check if the child node is a leaf node $S_1$. If the child node is $S_1$, then the tree is complete. However, if the child node is not $S_1$, then check that child node for all possible child nodes. In other words, check for all the possible reactions that have this node as a product. Repeat this process for each child node until only leaf nodes are left, $S_1$. Figure made with BioRender.

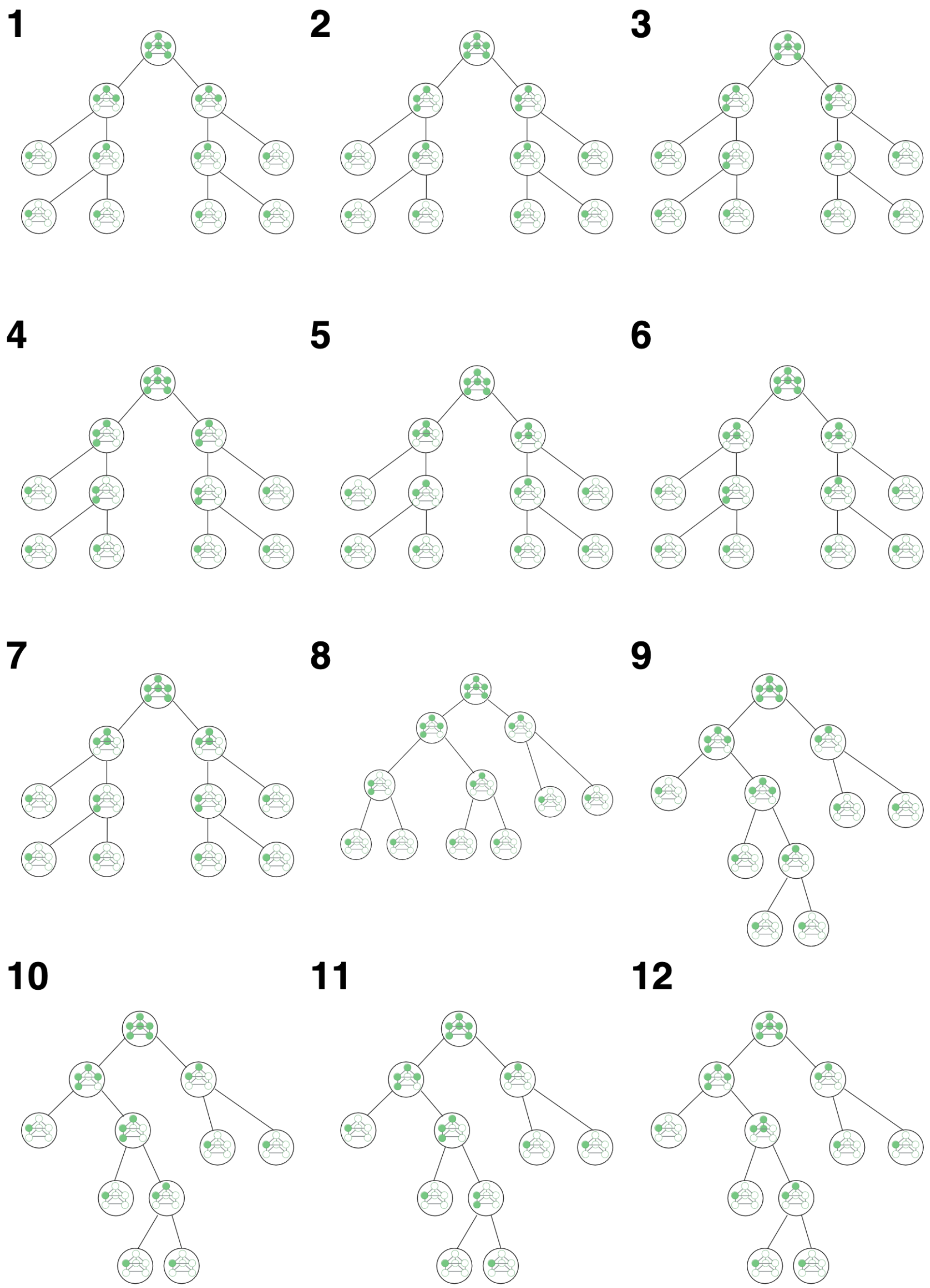

**Figure S12: All assembly pathways for the stacked trimer**. Each panel represents an assembly pathway for forming the stacked trimer. Each node is an intermediate species, we included the schematic for each species. The root node is the fully assembled stacked trimer $S_{12}$ and the leaf nodes are the monomer species $S_1$. Figure made with BioRender.

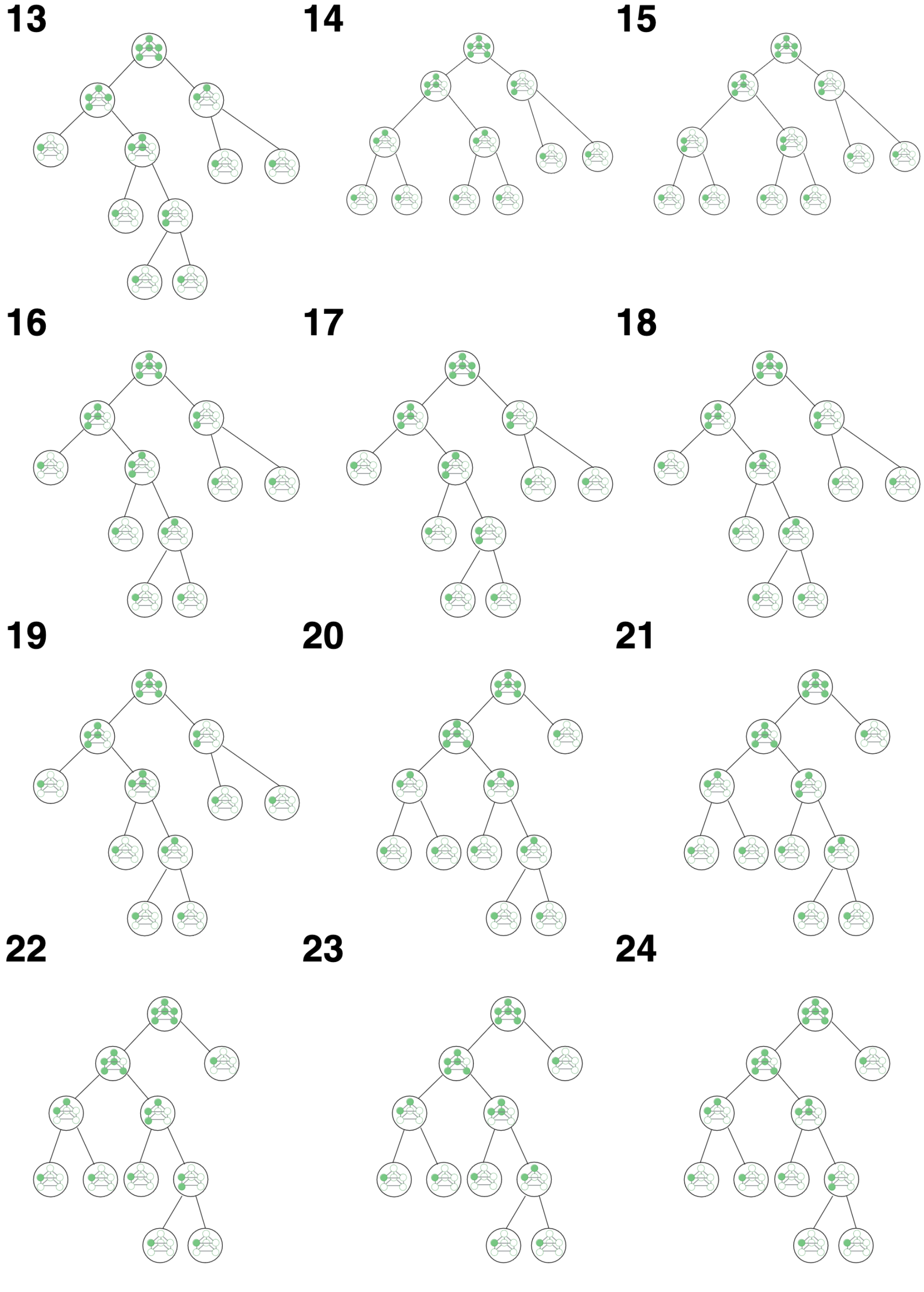

**Figure S13: All assembly pathways for the stacked trimer continued.**

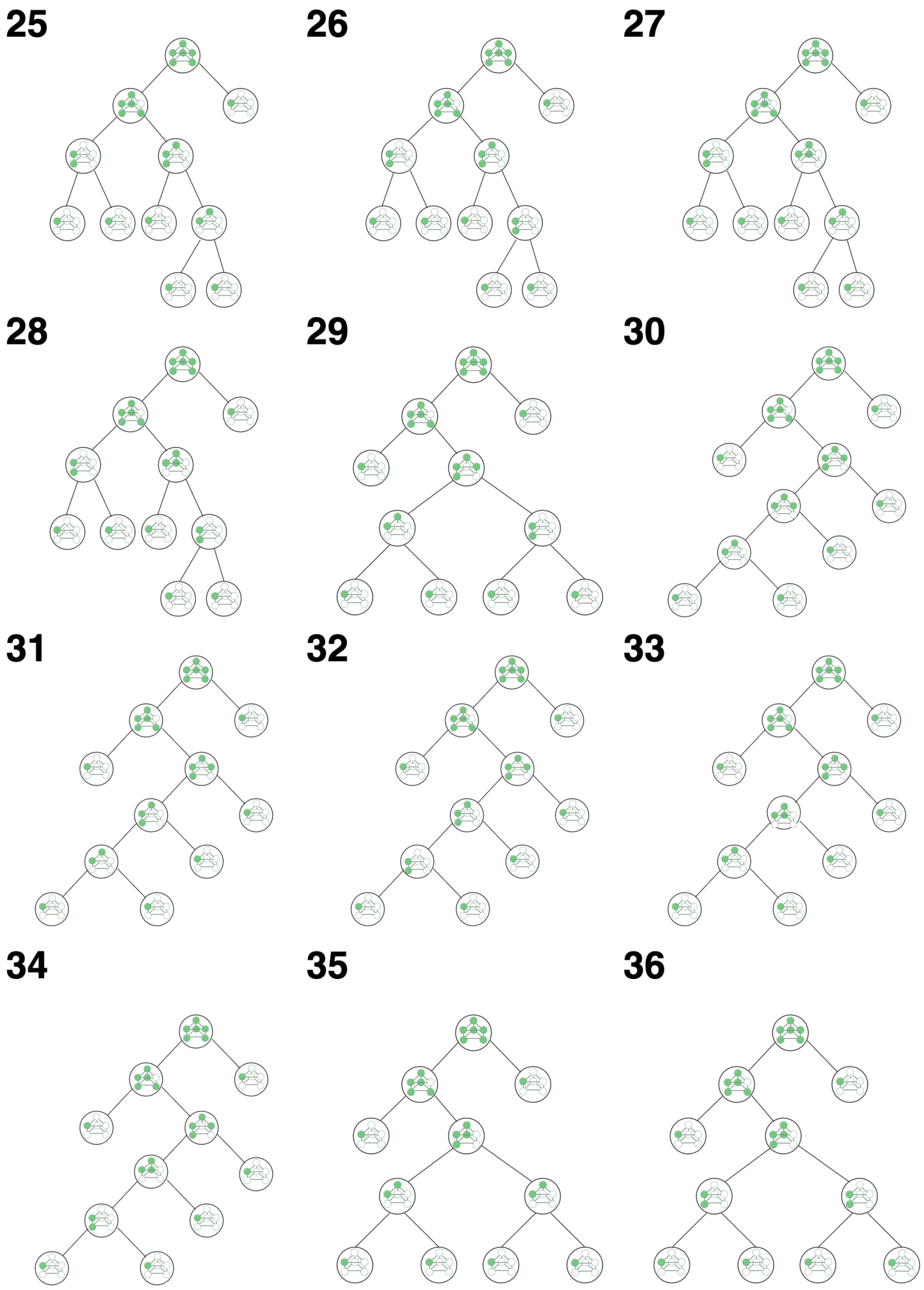

**Figure S14: All assembly pathways for the stacked trimer continued.**

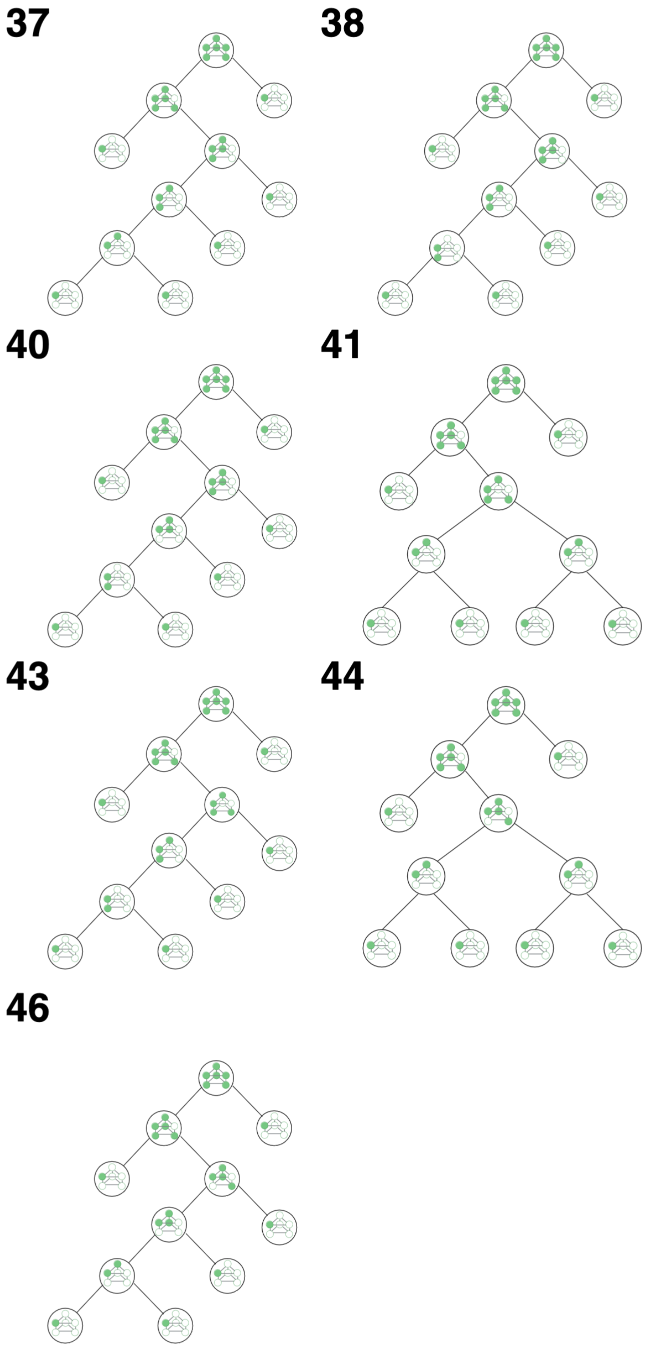

**Figure S15: All assembly pathways for the stacked trimer continued.**

For any species $S_i$, we can define the *total production flux* of that species as:

$$C_T(S_i) = \sum_{r_j \in \phi(S_i)} v_j(S_i)\mathcal{F}_{f,j}.$$

Note this is just the total amount of this particular molecule that is being made at any given time. While this value clearly varies with time, we keep that dependence implicit. We can now define the *relative contribution* of reaction $r_i$ to production of some species $S_j$ as:

$$C_{r_i}(S_j) = v_j(S_j)\mathcal{F}_{f,j}/C_T(S_j),$$

which is just the flux of production of $S_j$ by this reaction divided by the total flux of production of that species. Interestingly, this represents the *probability* that a molecule of $S_j$ that was produced in an infinitesimal time window at some time $t$ was produced *by* reaction $r_i$.

We now want to compute the contribution of a given pathway $P_k$ to the formation of the stacked trimer. By definition, the last "step" in any pathway is a reaction that forms the stacked trimer, $S_{12}$; in the example given in Figure S9, this is the reaction of $S_{11}$ with $S_1$. Call the reaction that produces the stacked trimer from these two species $r_i$. This last step will have a relative contribution $C_{r_i}(S_{12})$ as defined above. This represents the *total* contribution to $S_{12}$ formation by all pathways that have reaction $r_i$ as their last step.

To compute the contribution of a specific pathway, we have to consider not just the formation of the final structure $S_{12}$, but also the formation of the other intermediates in that pathway. Note that, in this example, we have $S_1$ reacting with $S_{11}$; since $S_1$ is the monomer, we need not consider how it is formed. There are, however, multiple reactions that can form $S_{11}$, and each of those reactions will correspond to different assembly pathways. Say we want to consider the contribution of pathways where $S_{11}$ is formed by the reaction of $S_8$ (the four-membered ring that forms the 'face' of the stacked trimer) and the monomer $S_1$. Let's call this reaction $r_k$. We can similarly calculate the relative contribution of reaction $r_k$ to the formation of $S_{11}$; this is just $C_{r_k}(S_{11})$.

Now consider the contribution of pathways that have *both* $S_1 + S_{11} \longrightarrow S_{12}$ as their last step *and* stipulate that $S_{11}$ was formed by the reaction $S_1 + S_8 \longrightarrow S_{11}$. This is just the probability that the last step is $r_i$ and the second-to-last step is $r_k$. So we can multiply the two probabilities together ($C_{r_i}(S_{12})$ for the first reaction and $C_{r_k}(S_{11})$ for the second reaction).

We can recursively apply this logic to each binary step in the pathway, and realize that the pathway contribution is just the product of all these relative fluxes. Thus, the relative contribution of an entire pathway $P_k$ is:

$$C(P_k) = \prod_{r_i \in P_k} C_{r_i}(S_j),$$

where $r_i \in P_k$ is just indicating the set of all reactions in the pathway and $S_j$ as the product of reaction $r_i$ is left implicit. By the definition of $C_{r_i}(S_j)$ this is inherently normalized, so that:

$$\sum_{P_i \in \mathcal{P}} C(P_i) = 1,$$

where $\mathcal{P}$ is the set of all pathways. In other words the sum over all pathway contributions is 1.

Note that, in the above case, we are implicitly considering the *in vitro* model, since we do not consider a reaction that creates the monomer species $S_1$. In the case of the *in vivo* model, there is technically a synthesis reaction that creates $S_1$, so this raises the question of how to extend our definition of pathway contributions to the *in vivo* case. Interestingly, there is only ever one reaction that can create the monomer, and so its pathway contribution to the formation will always be identically 1. Since there is only one way to form the monomer, neither the enumeration of pathways, nor the calculation of their contributions, is modified in that case. The only interesting difference for the *in vivo* case is that we can calculate the contribution of various pathways at steady-state. The results for pathway contributions shown in the main text are thus taken at time time $t = 8.64 \times 10^4$ s (i.e. 24 hours) in the *in vitro* case and at steady-state for the *in vivo* case.

#### REFERENCES

[1] Ramzi Alsallaq and Huan Xiang Zhou. "Electrostatic rate enhancement and transient complex of protein-protein association". In: *Proteins: Structure, Function and Genetics* 71 (1 Apr. 2008), pp. 320–335. ISSN: 08873585. DOI: 10.1002/prot.21679.

[2] Jieming Chen, Nicholas Sawyer, and Lynne Regan. "Protein-protein interactions: General trends in the relationship between binding affinity and interfacial buried surface area". In: *Protein Science* 22 (4 Apr. 2013), pp. 510–515. ISSN: 09618368. DOI: 10.1002/pro.2230.

[3] Eric J. Deeds, John A. Bachman, and Walter Fontana. “Optimizing ring assembly reveals the strength of weak interactions”. In: *Proceedings of the National Academy of Sciences of the United States of America* 109 (7 2012), pp. 2348–2353. ISSN: 00278424. DOI: 10.1073/pnas.1113095109.

[4] Anja Grziwa et al. “Dissociation and reconstitution of the Thermoplasma proteasome”. In: *Eur. J. Biochem* 223 (1994), pp. 1061–1067.

[5] Nancy Horton and Mitchell Lewis. “Calculation of the free energy of association for protein complexes”. In: *Protein Science* 1 (1 1992), pp. 169–181. ISSN: 1469896X. DOI: 10.1002/pro.5560010117.

[6] S. Mangan, A. Zaslaver, and U. Alon. “The coherent feedforward loop serves as a sign-sensitive delay element in transcription networks”. In: *Journal of Molecular Biology* 334 (2 Nov. 2003), pp. 197–204. ISSN: 00222836. DOI: 10.1016/j.jmb.2003.09.049.

[7] António J. Marques et al. “Catalytic mechanism and assembly of the proteasome”. In: *Chemical Reviews* 109 (4 Apr. 2009), pp. 1509–1536. ISSN: 00092665. DOI: 10.1021/cr8004857.

[8] David D.L. Minh et al. “The entropic cost of protein-protein association: A case study on acetylcholinesterase binding to fasciculin-2”. In: *Biophysical Journal* 89 (4 2005). ISSN: 00063495. DOI: 10.1529/biophysj.105.069336.

[9] Shigeo Murata, Hideki Yashiroda, and Keiji Tanaka. “Molecular mechanisms of proteasome assembly”. In: *Nature Reviews Molecular Cell Biology* 10 (2 Feb. 2009), pp. 104–115. ISSN: 14710072. DOI: 10.1038/nrm2630.

[10] Maulik K. Nariya et al. “Robustness and the evolution of length control strategies in the T3SS and flagellar hook”. In: *Biophysical Journal* 120 (17 Sept. 2021), pp. 3820–3830. ISSN: 15420086. DOI: 10.1016/j.bpj.2021.05.032.

[11] Hung D. Nguyen, Vijay S. Reddy, and Charles L. Brooks. “Deciphering the kinetic mechanism of spontaneous self-assembly of icosahedral capsids”. In: *Nano Letters* 7 (2 Feb. 2007), pp. 338–344. ISSN: 15306984. DOI: 10.1021/nl062449h.

[12] Xiaodong Pang, Sanbo Qin, and Huan Xiang Zhou. “Rationalizing 5000-fold differences in receptor-binding rate constants of four cytokines”. In: *Biophysical Journal* 101 (5 Sept. 2011), pp. 1175–1183. ISSN: 00063495. DOI: 10.1016/j.bpj.2011.06.056.

[13] Sanbo Qin, Xiaodong Pang, and Huan Xiang Zhou. “Automated prediction of protein association rate constants”. In: *Structure* 19 (12 Dec. 2011), pp. 1744–1751. ISSN: 09692126. DOI: 10.1016/j.str.2011.10.015.

[14] Leonor Saiz and Jose M.G. Vilar. “Stochastic dynamics of macromolecular-assembly networks”. In: *Molecular Systems Biology* 2 (May 2006). ISSN: 17444292. DOI: 10.1038/msb4100061.

[15] Ilya A Vakser et al. “Docking-based long timescale simulation of cell-size protein systems at atomic resolution”. In: (2022). DOI: 10.1073/pnas. URL: https://doi.org/10.1073/pnas.2210249119.

[16] Ilya A. Vakser and Eric J. Deeds. “Computational approaches to macromolecular interactions in the cell”. In: *Current Opinion in Structural Biology* 55 (Apr. 2019), pp. 59–65. ISSN: 1879033X. DOI: 10.1016/j.sbi.2019.03.012.

[17] Huan Xiang Zhou. “Rate theories for biologists”. In: *Quarterly Reviews of Biophysics* 43 (2 May 2010), pp. 219–293. ISSN: 00335835. DOI: 10.1017/S0033583510000120.
